# Development of human iPSC-derived cardiomyocytes-based heart-on-a-chip with customized multi-electrode array for drug-induced cardiotoxicity evaluations

**DOI:** 10.1101/2025.09.16.676208

**Authors:** Geonho Choi, Ki-suk Kim, Han-Yeol Yang, Sungwoo Cho, Song-ah Park, Dohyung Kwon, Si-hun Cho, Seunghyun Yoon, Hyun-soo Kang, Jae-hyeok Lee, Sungho Ko

## Abstract

The heart is an essential organ at the beginning and end of life. Cardiotoxicity directly related to life is emphasized as an absolute safety factor in drug development, so it is indispensable to evaluate it accurately. That is why establishing a novel platform that can precisely recapitulate human cardiophysiological properties is necessary. Here, we report a human induced pluripotent stem cell (hiPSC)-derived cardiomyocytes-based heart-on-a-chip integrated with a customized multi-electrode array (cMEA), capable of actualizing a cardiac muscle fascicle and surrounding capillaries tridimensionally. The differences in experimental reproducibility, scale of field potential wave, and gene expression patterns according to the two-dimensional cell-level and three-dimensional tissue-level culture are confirmed. Excellent accuracy of drug efficacy/toxicity of the chip is also verified relative to other conventional cell culture platforms by looking at heart rate changes depending on drug type and concentration/time. Additionally, we checked that kinetic/electrophysiological/molecular biological analysis can be performed with our chip by applying motion tracking, cMEA, and RNA sequencing. Thus, our platform is expected to be used as a tool for more exact cardiotoxicity tests with high reproducibility and multi-angle analysis in line with new pharmacological safety assessments such as comprehensive cardiac arrhythmias assessment (CiPA).

## 1. Introduction

The heart is an essential organ that maintains life by continuously supplying nutrients and oxygen to every corner of the body and removing wastes and carbon dioxide through blood circulation. However, due to these constantly moving features, various diseases occur in the heart if there is even a little abnormality. Unlike other diseases, cardiac disorders can directly lead to sudden death, so it is urgent and vital to establish safety pharmacology assessments for cardiac side effects. The competitiveness of newly discovered drugs depends on ‘safety,’ and cardiotoxicity is particularly one of the most important safety factors. Since some medicines were withdrawn from the market due to Torsades de Pointes (TdP), a symptom of fatal arrhythmia, a large portion of the withdrawal drugs over the past years caused heart disorders have been found [1–3]. Although diverse studies on heart diseases caused by drug toxicity are currently in progress, the incidence and mortality rates worldwide are continuously increasing. Thus, it is clear that it is time for more definitive and in-depth research on heart disease-drug interactions.

Currently, assessment of drug-induced cardiotoxicity for drug development is exclusively conducted using an hERG assay. The hERG assay, one of the non-clinical evaluation methods to confirm the prolongation of ventricular repolarization, can test whether a drug or its metabolites induce hERG block [4]. Nonetheless, it has recently been recognized that cardiotoxicity evaluation with hERG assay alone is somewhat inaccurate as false negatives may occur. Therefore, the need to present a new paradigm for more accurate cardiotoxicity evaluation has emerged. In 2013, the International Council for Harmonization of Technical Requirements for Pharmaceuticals for Human Use (ICH) established a project group called the comprehensive in vitro pro arrhythmia assay (CiPA) initiative to seek novel cardiac safety pharmacology tests. CiPA comprehensively tests major channels of cardiac action potentials that affect arrhythmias, such as sodium and calcium channels as well as hERG channels, improves the accuracy of drug cardiotoxicity prediction through in silico computer modeling, proposes human induced pluripotent stem cell-derived cardiomyocyte (hiPSC-CM) based in vitro action potential analysis, and suggests the need for new biomarkers along with existing indicators [5–7].

Among the contents proposed in CiPA, hiPSC-CM-based in vitro action potential analysis will be particularly suitable to be performed through heart-on-a-chip. Animal testing, an existing toxicity evaluation model, is a global trend regulated due to ethical issues such as animal welfare and inaccuracies due to species specificity. At present, developed countries have established centers for alternatives to animal testing [8–9] and prepared policies for regulating animal testing, such as the case that the US Environmental Protection Agency (EPA) announced that they will provide support to prohibit all animal testing by 2035 [10–11]. Thus, the toxicological assessment technology using animals is expected to disappear gradually, mainly in developed countries, and human-like modelings based on hiPSC-CMs will be utilized [6]. Using hiPSC-CMs is expected to reduce animal usage and improve mechanism-oriented understanding of changes in TdP or cardiac action potential based on human genetic background. Besides, the various in vitro action potential analyses proposed by CiPA include a multi-electrode assay (MEA). The MEA system is a technology that detects and records changes in electrical activity, such as the emission or reception of intercellular signals, through approximately 60 arrayed multi-electrodes. This technology can be measured while cells are alive and monitor cell responses in real-time while applying external stimuli such as chemical or physical stimuli. Thus, measuring extracellular field potentials generated by cardiomyocyte microcurrents can be utilized as parameters for evaluating the ventricular repolarization delay induced by drugs. Measured values such as field potential duration (FPD), beat per minute (bpm), interspike interval (ISI), and dv/dtmax are representative [12–14]. Hence, there is a need to actively prepare for the international trend regarding the ICH cardiotoxicity assessment guidelines by reflecting the usage of the hiPSC-CM and MEA systems proposed by the CiPA in research.

Heart-on-a-chip research has been studied in three representative forms. First, the thin film-based chip design, which can observe heartbeats via the phenomenon that the cardiomyocytes-cultured thin film is rolled up by cell contraction, has been improved to date and applied to various related studies [15–23]. However, this design is difficult to call a three-dimensional (3D) culture, as cells are cultured on thin films two-dimensionally (2D). Most of them are in the form of wells that are no different from conventional culture-ware, so fluid flow is not applied. Moreover, heartbeats are mechanically measured according to the shape change of the film, so electrophysiological analysis seems difficult. Second, the chip designs using micro-pillars, or cantilevers, are generally called a term of engineered heart tissue (EHT), which is also included in the category of heart-on-a-chip [24–30]. When cardiomyocytes are cultured near two micro-pillars, they are spontaneously aligned and organized along the axis between the mico-pillars, forming the most matured myocardial tissue among all chip designs. Nonetheless, this is also in the well-type, so there is no fluid flow. Electrophysiological analysis is similarly not feasible since heartbeats are mechanically gauged based on the degree of curvature of the pillars. Lastly, the chip design with microfluidic structures selected by most research groups can be 3D cultured and fluid flow is applied [31–39]. Although there are studies that have analyzed heartbeats using calcium imaging or electrophysiological measurements, they are rare, and no instances of mechanical measurement have been identified. Recently, research cases have been reported that combine the advantages of these three designs [40–44]. Additionally, heart-on-a-chip studies incorporating MEA have also been reported [45–49]. However, most of them conducted 2D culture despite microfluidic structures or applied well-type for culturing cells on electrodes like regular MEAs.

Herein, we developed a human stem cell-based heart-on-a-chip integrated with a customized multi-electrode array (cMEA) to recapitulate cardiac muscle fascicle and surrounding capillaries and monitor exact drug responses. To overcome the limitations of inter-species differences in animal experiments and to replace the cancer-derived cell lines with different physiological activities, we used hiPSC-CMs based on the human genome. We devised a new micro-post structure, applied silicone adhesive to chip fabrication, and customized multi-electrodes to the heart chip to facilitate our experiment. The superiority of the 3D tissue-level culture using collagen, the extracellular matrix (ECM) of actual myocardial tissue, was confirmed compared to the 2D cell-level culture, which is distinct from the in vivo microenvironment. We treated isoproterenol and nifedipine as the representative drugs that affect heart rate and observed the dose-dependent and time-lapse changes using kinetic, electrophysiological, and molecular biological analyses by applying motion tracking, cMEA, and total RNA-sequencing. Consequently, using our chips, it was possible to compare the differences in drug response according to the 2D and 3D cultures and to confirm the possibility of drug-induced cardiotoxicity evaluation. Our platform is the first heart-on-a-chip that realizes the perimysium structure at the chip, and it is expected to be a timely cardiotoxicity test platform that can be used for CiPA projects based on its high reproducibility and versatility capable of multi-angle analysis.

## 2. Results

### 2.1. Design of heart-on-a-chip

#### 2.1.1. Biomimetic points of heart-on-a-chip

To design a heart-on-a-chip that can recapitulate cardiophysiological features as much as possible, we sought significant biomimetic points of the heart that can be realistically actualized in the chip. First, looking at the overall histological microstructure of the heart, the most basic cardiac unit is cardiomyocytes, which are structurally connected to form myocardial fibers. The cardiac ECM, containing type 1 collagen as a principal constituent [50–55], shapes a collagen layer called the endomysium that swathes each cardiac muscle fiber. Also, these several myocardial fibers and endomysium are enwrapped by another thin collagen layer, the perimysium, to form a cardiac muscle fascicle [56]. Numbers of these myocardial fascicles gather to shape cardiac muscle tissue, the myocardium, and it does not have an additional collagen layer called the epimysium surrounding the myocardial fascicle bundles, unlike skeletal muscle tissue. This myocardium, together with the endocardium and epicardium, finally composes the heart wall. Additionally, capillary vessels branch off from coronary vessels and spread evenly into the myocardium to supply nutrients and oxygen and to discharge waste products and carbon dioxide. There are currently technical limitations in biomimicking all the functions of the heart into the chip, and the complexity may act as an obstacle to performing the drug efficacy/toxicity tests even if cardiac functions can be somewhat realized. Hence, we thought that the heart-on-a-chip should be designed to be reproducible with a simple structure and simultaneously include essential functions and structures of the heart. In consequence, among the structural ranges within the heart described above, the organized cardiac muscle fascicle containing the cardiomyocytes, endomysium, perimysium, and peripheral capillaries was selected as the appropriate biomimetic points (**Figure 1**).

**Figure 1.**
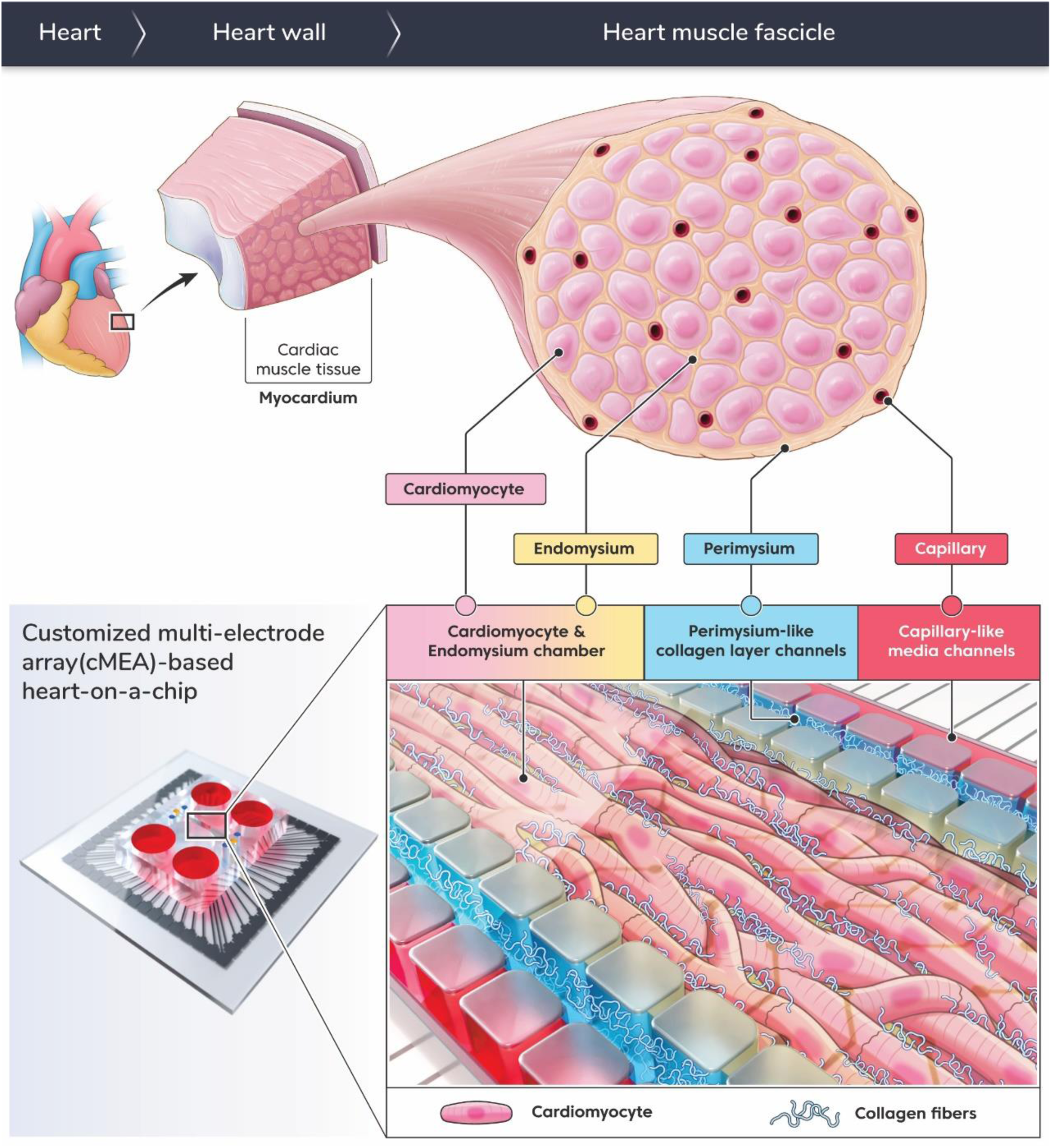
Schematic design of heart-on-a-chip integrated with cMEA. This chip recapitulates a cardiac muscle fascicle, which contains myocardial fibers composed of cardiomyocytes, collagen layers called endomysium and perimysium, and surrounding capillaries. The in-chip microstructures comprise a central cardiomyocyte and endomysium chamber, adjacent two perimysium-like collagen layer channels, two outermost capillary-like media channels, and four micro-post arrays at intervals among the chamber and the channels. All electrode probes are placed in the chamber.

#### 2.1.2. Schematic design of heart-on-a-chip

An ideal arrangement of microstructures inside the chip to establish in vitro culture conditions similar Therefore, the in-chip microstructures consisted of a central cardiomyocyte and endomysium chamber, two adjacent perimysium-like collagen layer channels, two outermost capillary-like media channels, and four micro-post arrays at the spaces between the chamber and channels. This arrangement allowed the media flow from the media channels to pass through the collagen layer channels and finally to the cell chamber, just as nutrients and drugs are delivered to cardiomyocytes from capillaries across collagen layers like perimysium and endomysium. Also, the media channels were placed on the outermost side to minimize shear stress directly given to the cardiomyocytes, and the diffusion-like media flow was simultaneously induced between the gaps of collagen fibers by forming 3D collagen layers between the cell chamber and media channels [57–58]. In addition, the width of the cardiomyocyte and endomysium chamber was set to 600 μm based on the diameter of the myocardial fascicle in vivo of 150-200 μm [36, 56], the width of the perimysium-like collagen layer channel was set to 60 μm on the basis of the actual perimysium diameter of 20 μm [59], and the width of the capillary-like media channel was set to 30 μm with reference to the actual capillary diameter of 10 μm [60–63]. On this wise, all in-chip microstructures were set to magnificate about three times the actual size for reproducible chip fabrication, taking account of the in vivo microstructure.

A micro-weir structure in an arc form with a thickness of 40 μm in the cardiomyocyte and endomysium chamber just before the narrowing to the outlet was located. We hereby maintained a constant cell density in the cell chamber by preventing cell loss to the outlet as much as possible, leading to securing experimental reproducibility. Additionally, this micro-weir, which is 10 μm high from the chip bottom, allowed the cell density to be ideally controlled by applying a suction force to the outlet side to trap the cells in the micro-weir, whereas to selectively drain the collagen solution into 10 μm space below the structure. The aforementioned micro-post structures arranged in the four rows were devised in an asymmetrically rounded rectangular shape, and they were arranged between the chamber and the channels so that all fluids injected into the chip could be separately infused into the desired space without mixing. The size of these micro-posts was set to 120 μm width x 100 μm depth, and the distance between each micro-post was 20 μm. The height of all microstructures inside the chip was set to 50 μm, except for the micro-weir mentioned above.

The multi-electrode array custom-made for the heart-on-a-chip, cMEA, was set to 49 mm width x 49 mm depth x 1 mm height to be compatible with a head stage of the MEA2100-system (Multichannel systems), which is the component of the existing measuring equipment for MEA connection. The total electrode of the cMEA was composed of 59 recording electrodes and 1 reference electrode by referring to the standard MEA design, but the entire electrode grid arrangement was arranged in 4 x 15 so that all electrodes could be positioned in the cardiomyocyte and endomysium chamber, and the distance between each recording electrode was placed 150 μm apart. The diameter of the recording electrodes is 30 μm, the diameter of the reference electrode is 50 μm, the diameter of the track connecting them is 10 μm, and the contact pad connected to the equipment is 2.2 mm width x 2.2 mm depth was set.

Conclusionally, we reflected the anatomical/physiological characteristics of the cardiac muscle fascicle and surrounding capillaries, which are the structural morphological ranges within the heart, to realize accurate cardiophysiological phenomena in the chip. As for the experimental aspect, structural elements such as micro-weirs and micro-posts were added to obtain more reproducible results through simple manipulation. Finally, we enabled more precise cardiotoxicity tests by integrating the cMEA for functional analysis of myocardial tissue to be actualized inside the chip (**Figure 1**) (**Movie S1**).

### 2.2. Characterization of heart-on-a-chip

#### 2.2.1. Fluid injection without mixing

Characterization of the fabricated heart-on-a-chip in this research was successfully performed (**Figure 2**). In-chip microstructures capable of isolating the fluids are generally required for fluid injection into the adjacent spaces without mixing. Micro-valves are applied in most cases but require complex microstructures, relatively cumbersome operation, and additional external driving force [64–65]. Trapezoidal micro-post arrays are also widely utilized, but this shape is vulnerable to fluid intermixing in the adjoining space and is poorly reproducible, even if successful [57]. Hence, we devised asymmetrically rectangular micro-posts with rounded corners arranged in a zigzag pattern similar to a zipper to improve these aspects. Firstly, the reason why the round corners were adopted was that these were to shape bottlenecks by narrowing the space between the micro-posts so that surface tension between the micro-posts could be maximized. In addition, each boundary of the injected fluid was formed at the narrowest parts of the bottlenecks so that pneumatic gaps were naturally created between the far-distant fluid borders to isolate contiguous spaces without complex microstructures in a simple way. The asymmetric circular edges of the micro-posts with small radii toward the perimysium-like collagen layer channels allowed the fluid boundaries to be located as close as possible to the channels so that the thickness of the collagen layer was not partially thickened. On the other hand, the bottleneck effect was minimized due to the relatively small radii, so it was challenging to maintain the surface tension between the micro-posts against the injecting pressure. Besides, it was very tricky to separately infuse viscous fluid such as collagen solution, which is not general fluid. To get around this problem, we introduced the zigzag pattern arrangement by contemplating two physical phenomena: i) In fluid statics, meniscus is a phenomenon in which the upper surface of liquid in a solid container becomes curved in a convex or concave form along the boundary of the container by surface tension. In the case of water-based liquids, the adhesive force between liquid and solid is greater than the cohesive force between liquid molecules [66], so we thought the water-based collagen solution would move to rely on the solid micro-posts. ii) Another phenomenon is that the interfacial tension between liquid and solid is usually weaker than the surface tension between liquid and gas [67], so we guessed that the injecting pressure would advance in the infusing direction, which is the liquid-solid interface, rather than in the direction between the micro-posts, which is the liquid-gas interface. High-temperature treatment was carried out in order to reinforce the surface tension inside each chip further, thereby securing experimental reproducibility [68]. Lastly, we injected food coloring to confirm that fluids could be selectively infused into the desired space only with a new type and arrangement of micro-posts mentioned above. Consequently, we checked that the gaps were well formed in the spaces between the micro-posts so that food coloring was not mixed, and it was possible to separate injection into the desired space. Moreover, the expected fluid movement by the zigzag pattern and the collagen layer formation with reproducibility were verified (**Figure 2a**) (**Movie S2**). Yet, we interpreted that the fluid was injected in a convex rather than a concave form because the infusing pressure was constantly applied rather than in a static situation, unlike the referenced phenomenon.

**Figure 2.**
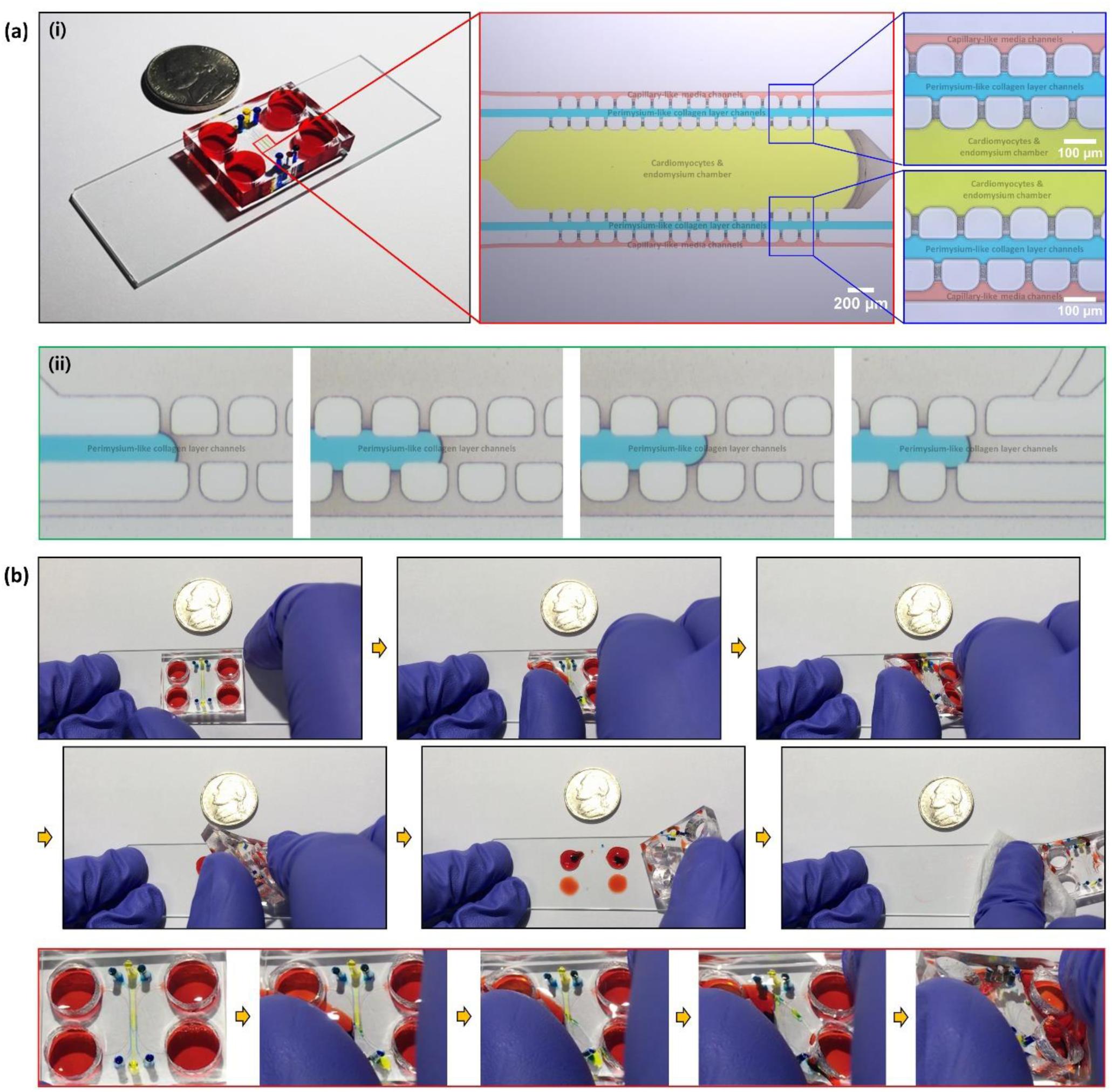
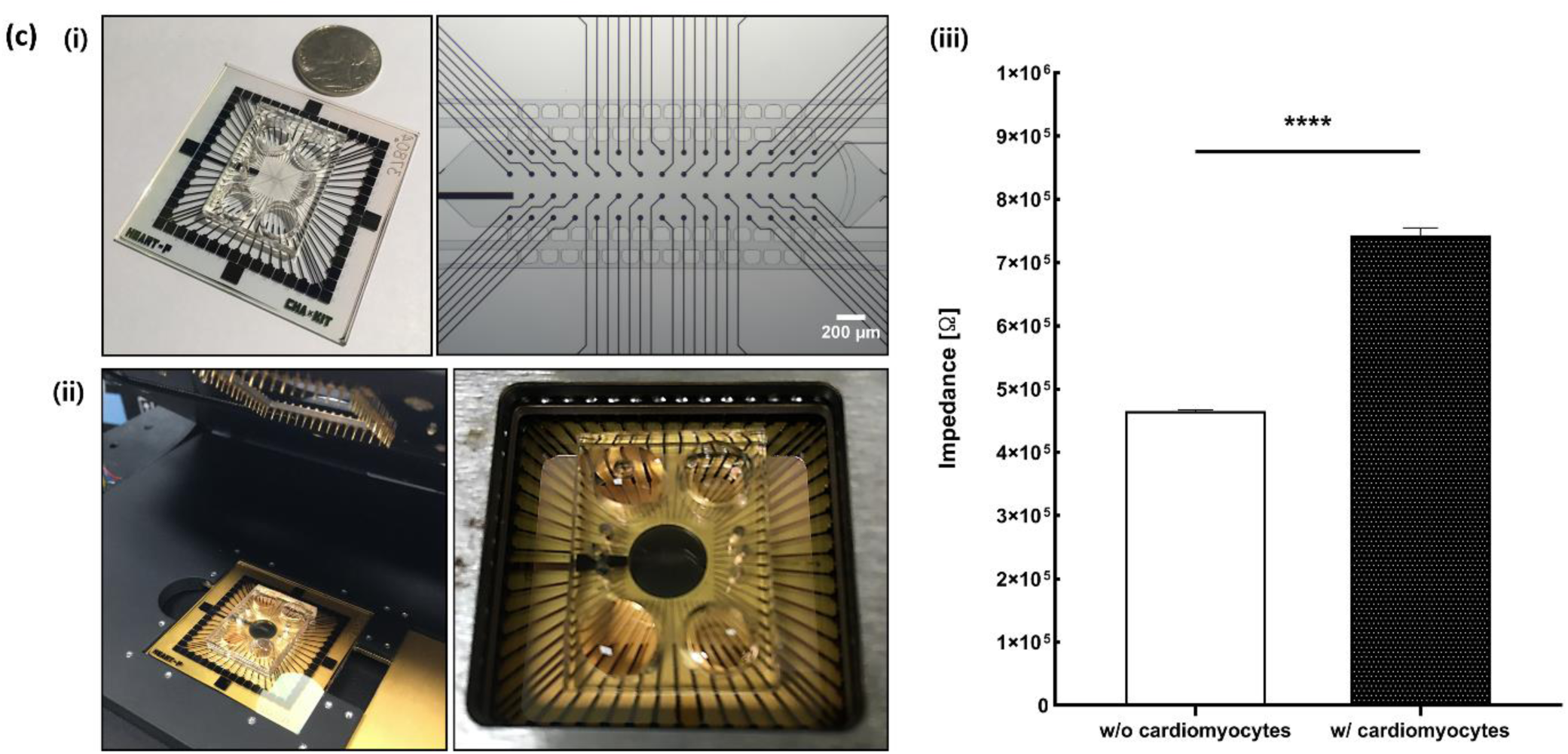
Characterization of the heart-on-a-chip. (a) Fluid injection without mixing. Zipper-like zigzag arrangement of asymmetrically rounded rectangular micro-posts could form pneumatic gaps at the interspaces between the micro-post arrays, enabling reproducible and straightforward separation infusion. Scale bars: 100-200 μm. (b) Reversible bonding of microfluidic units fabricated by PDMS and Silicone adhesive mixture. Silicone adhesive application for fabrication of microfluidic units could easily harvest cell tissue in the chip and reuse expensive electrode units like MEA. (c) Electrochemical characterization of cMEA. The cMEA shows great sensitivity to impedance changes according to the presence or absence of cardiomyocytes. Scale bars: 200 μm. Statistical analysis was performed via unpaired t-test. \*\*\*\**P* <0.0001. All data are mean ± SD.

#### 2.2.2. Reversible bonding of microfluidic units and glass substrates

Reversible bonding was a prerequisite that we must be established to conduct our experiment more efficiently. The adhesion between the polydimethylsiloxane (PDMS)-based microfluidic units and the glass substrates by the well-known oxygen plasma treatment is irreversible bonding, which has excellent adherence. However, on the contrary, this makes it challenging to harvest cultured cell tissues from the chips for molecular biological analysis without damage, and their utility is limited because they cannot be reused when expensive electrode units such as MEA are applied. Therefore, we had to consider a reversible bonding method that does not affect the existing research results, retains the attaching force without fluid leakage even after long-term culture, can recover cell tissue in the chip, and does not generate marks or fragments during detachment for reuse. In previous studies, several reversible bonding means that were resetting the existing oxygen plasma conditions to weaken sticking or utilizing coating materials such as dimethyl-methylphenylmethoxy siloxane (DMPMS), poly l-lysine (PLL), and gelatin have been proposed [69–72]. We particularly paid attention to the silicone-based material used as a medical adhesive that the Chu M research group (2017) applied to reversible bonding [73]. The methods other than silicone adhesive were relatively hard to maintain adhesion, left marks, and required additional surface treatment, which was cumbersome. Besides, there was a possibility that surface denaturation of the MEA substrates could occur, causing variables in measurements using them. On the other hand, the silicone adhesive was contemplated because of satisfactory adhesion, no marks when detached, and no surface treatment with its own adhesive strength was required. Moreover, it was biocompatible, as it was used for medical purposes, and easy to apply because it was the same silicone-based material as PDMS.

However, due to the discontinuation of the silicone adhesive products used in previous studies, it was necessary to establish a protocol considering the physical properties of the newly refurbished materials. For reversible bonding, microfluidic units must be manufactured by mixing previously used PDMS with a silicone adhesive in an appropriate ratio. The proportion of the PDMS:silicone adhesive mixture was set to 1:3 by first considering the adhesion that can culture for a long period and the hardness that can punch reproducibly. As mentioned above, the portion of silicone adhesive, which is relatively weak in stiffness, became proportionally dominant when mixing. Thus, the solidity of the cured PDMS:silicone adhesive mixture became soft as well, thereby making it difficult to punch inlets, outlets, and media reservoirs during the manufacturing process. On account of this problem, the ratio of PDMS:curing agent is usually set to 10:1, but we modified it to 8:1 to balance by increasing the rigidity of the PDMS part. Before cell culture, we checked whether the desired level of reversible bonding was possible by applying silicone adhesive after injecting food coloring with a syringe pump. As a result, we verified that there was no effect on separation infusion because fluid mixing in the chip did not occur even with the application of silicone adhesive. It was also confirmed that it came off cleanly without leaving any marks or fragments (**Figure 2b**) (**Movie S3**).

#### 2.2.3. Electrochemical characterization of cMEA

We evaluated whether the newly manufactured multi-electrode array customized to the heart-on-a-chip functions appropriately through impedance change depending on the presence or absence of cells preliminary to the electrophysiological measurement of the cardiomyocytes. For this, the impedance differences between the condition cultured cardiomyocytes on the chamber of the heart-on-a-chip integrated with the cMEA and the condition loaded only media without cells were comparatively analyzed. As a result, there was no abnormality in the sensitivity of the cMEA by verifying that the impedance value significantly increased on account of the cardiomyocytes on the microelectrodes of the cMEA (**Figure 2c**, w/o cardiomyocytes 4.65 x 10^5^ Ω ± 0.02; w/ cardiomyocytes 7.43 x 10^5^ Ω ±0.11).

### 2.3. Culture of hiPSC-CMs on the heart-on-a-chip

There were several experimental considerations to establish a protocol for ideally infusing/culturing hiPSC-CMs at the heart-on-a-chip (**Figure 3a**). When injecting the collagen-based hydrogel solution into the perimysium-like collagen layer channels, it was necessary to proceed only at a low speed of about 3-5 μl/min. This was because the pressure generated by high-speed infusion not only intruded the fluid toward neighboring chambers and channels but also raised a problem in reproducible collagen layer formation. We thought that the cause of this phenomenon was because the collagen solution was more viscous than general solutions, so the injecting pressure was relatively intense, easily breaking down the surface tension between the micro-posts. In addition, even if a desired linear type of collagen layer was formed at the channels, the collagen fibers in the solution were not evenly spread, and most were swept out through the outlets connected to the channels through the infusing pressure. This gave rise to weakening the hardness of the collagen layer or maintaining a liquid state without morphological change, even if gelation was performed later. As another consideration, we had to place the collagen solution in ice for at least 1 h as the low-temperature preparation process before infusing it into the collagen layer channel. Sung KE Research Group (2009) reported that preincubation at a low temperature below 4 °C strengthened the thickness of collagen fibers and increased cell viability [74]. Thus, in consideration of the temperature-based gelation of the collagen, it was chilled at a lower temperature of 4 °C and shifted at 37 °C to induce more solid gelation via drastic temperature change. In consequence, in the low-speed infusion and the low-temperature conditions, we confirmed that the collagen fibers were more evenly distributed and distinctly observed, and the collagen layers also represented constant hardness with reproducibility (**Figure 3b**). Finally, we had to carry out two gelation steps by dividing each injection step of the collagen solution and the cell-collagen mixture rather than simultaneously infusing them and conducting only one gelation. When only one gelation was executed at 37 °C after the simultaneous injection, the temporarily formed gaps between the micro-posts collapsed, causing cell loss into the collagen layer that was not entirely gelated yet. Therefore, by subdividing these steps, we preceded the collagen layer gelation to indurate it so that cell loss could be prevented even if the gaps collapse on account of temperature changes (**Figure 3b**) (**Movie S4**). After all infusions, we checked that the gaps temporarily formed disappeared naturally at room temperature, and the ideally cultured 3D myocardial tissue at the heart-on-a-chip has a heart rate within 60-90 beats, which is known as the normal range (**Movie S5**). Additionally, we checked whether cardiomyocytes retain their identity well despite in-chip culture through all positive signals of actin cardiac muscle 1 (ACTC1), actinin alpha 2 (ACTN2), connexin 43 (Cx43), cardiac troponin T (cTnT2), which are typical cardiac markers (**Figure 3c**). Thus, we could be convinced that the heart-on-a-chip is suitable for culturing healthy myocardial tissue and is ready for cardiotoxicity evaluation.

**Figure 3.**
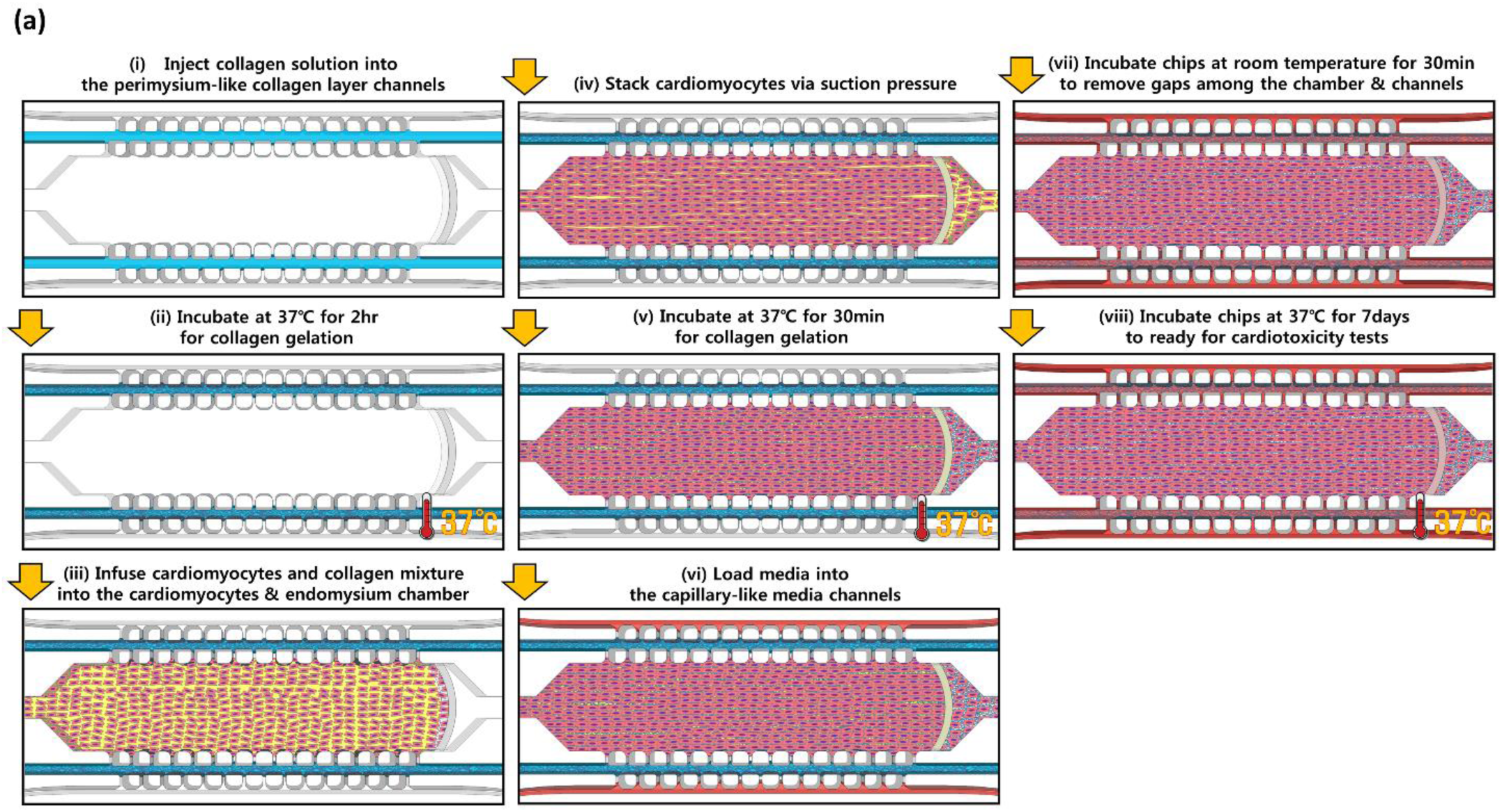
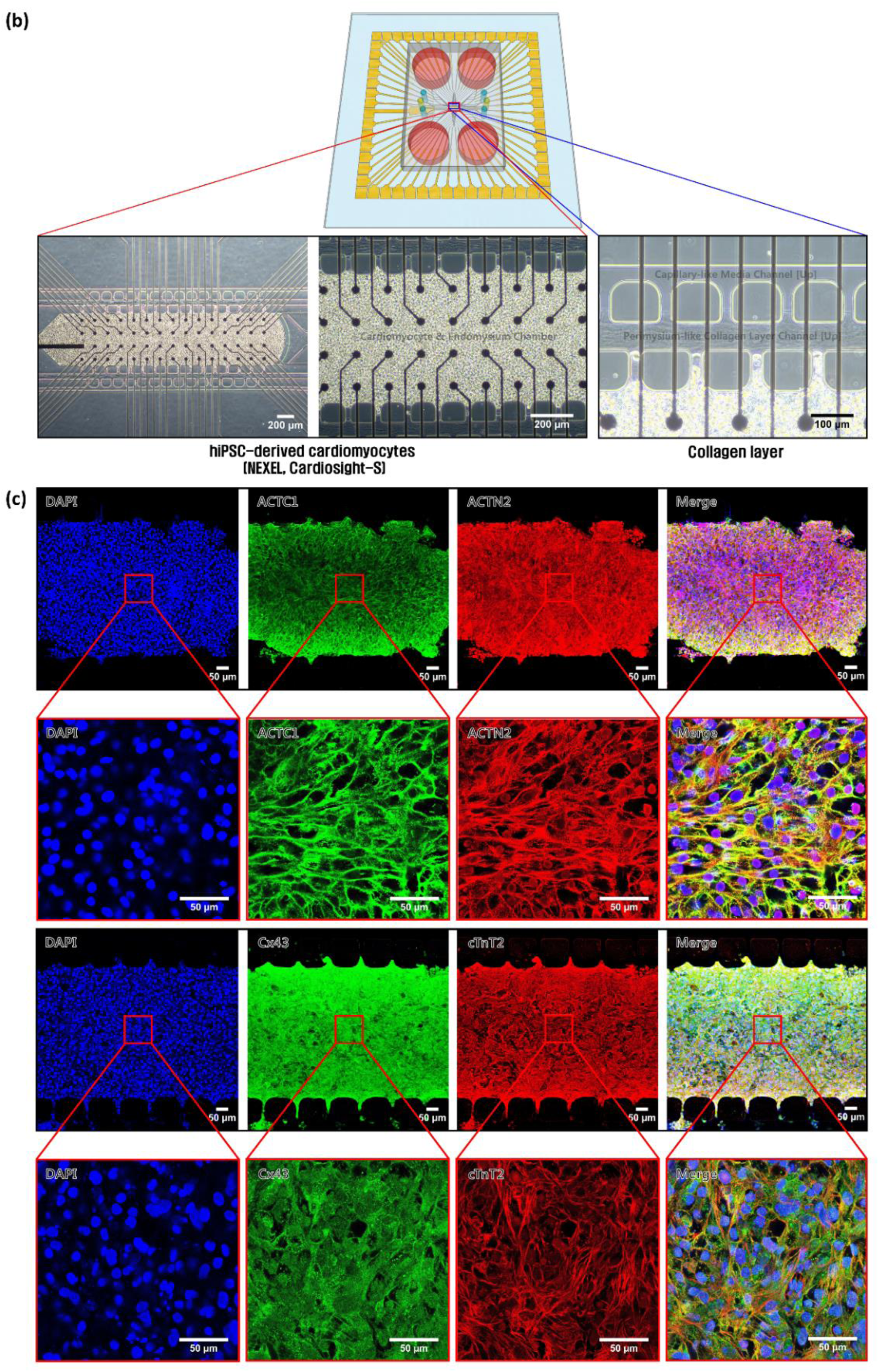
Culture of hiPSC-CMs on the heart-on-a-chip. (a) Flow chart for cell loading and culture. (b) Injection of hiPSC-CMs into the heart-on-a-chip. Scale bars: 100-200 μm. (c) Immunocytochemistry of hiPSC-CMs on the heart-on-a-chip (cardiomyocyte-specific markers, ACTC1 and Cx43, green; ACTN2 and cTnT2, red) (DAPI, blue). Scale bars: 50 μm. Figures 3(b) and 3(c) indicate that the hiPSC-CMs within the chip were maintained in a healthy state, and the cells retained their identity even inside the chip.

### 2.4. Comparative analysis of hiPSC-CMs on cell culture plate (2D) and heart-on-a-chip (3D)

#### 2.4.1. Difference in the heart rate during culture optimization

The optimal conditions for culturing hiPSC-CMs in the heart-on-a-chip were set, and at the same time, the significance of the chip was evaluated through a comparative analysis of 3D culture using the chip and the 2D culture using the conventional culture-ware. To this end, as the intuitive approach, we checked how long it took to culture until the hiPSC-CMs in all the chips reached within the range of 60-90 beats per minute (bpm), known as the normal heart rate. Whether reproducibility was secured for each chip was tested, and observation by 12 well plates was also performed. As a result, in the case of the heart-on-a-chip, no beating was observed in any of the chips on day 1 after injection, and the beating began to occur locally on day 2 and spread to the entire myocardial tissue on day 3, reaching an average of 56 bpm. On day 5, most chips gradually increased until an average of 64 bpm, and all chips came within the normal heart rate range on day 7 and kept an average beat rate of 65 bpm. Accordingly, we determined that 7 days was the optimal culture period within the chip and, at the same time, the optimal time to conduct drug efficacy/toxicity evaluation. The difference in heart rate between the chips was relatively uniform during the culture period, and reproducibility was also confirmed by observing that the gap decreased to around 5 bpm on day 7. On the other hand, in the case of the 12 well plates, an average of 68 bpm was observed from day 1 after injection and further increased on day 3 to an average of 98 bpm. On day 5, an average of 120 bpm was observed, exceeding the normal range, and on day 7, the heartbeat decreased again to an average of 96 bpm. The critical point is that the heart rate differed depending on the observation location, even within the same well, from day 2 to day 4. On day 5, the heartbeats within each well were synchronized, but the gap in heat rate between the wells was abnormally large, ranging from a maximum of 284 bpm to a minimum of 70 bpm. Since the difference in heart rate was severe even on day 7, as a temporary measure, we had no choice but to select only wells with a rate close to the normal range to evaluate drug efficacy/toxicity with equal comparison (**Figure 4a**) (**Movie S6, S7, S8, and S9**). Therefore, we judged that the existing 2D culture is unsuitable for accurate results because the experimental error was substantial due to the lack of reproducibility with the severe difference in heartbeats. On the contrary, we could prove that the 3D culture using cardiac biomimetic chips can obtain much more reproducible and accurate results than the existing methods.

**Figure 4.**
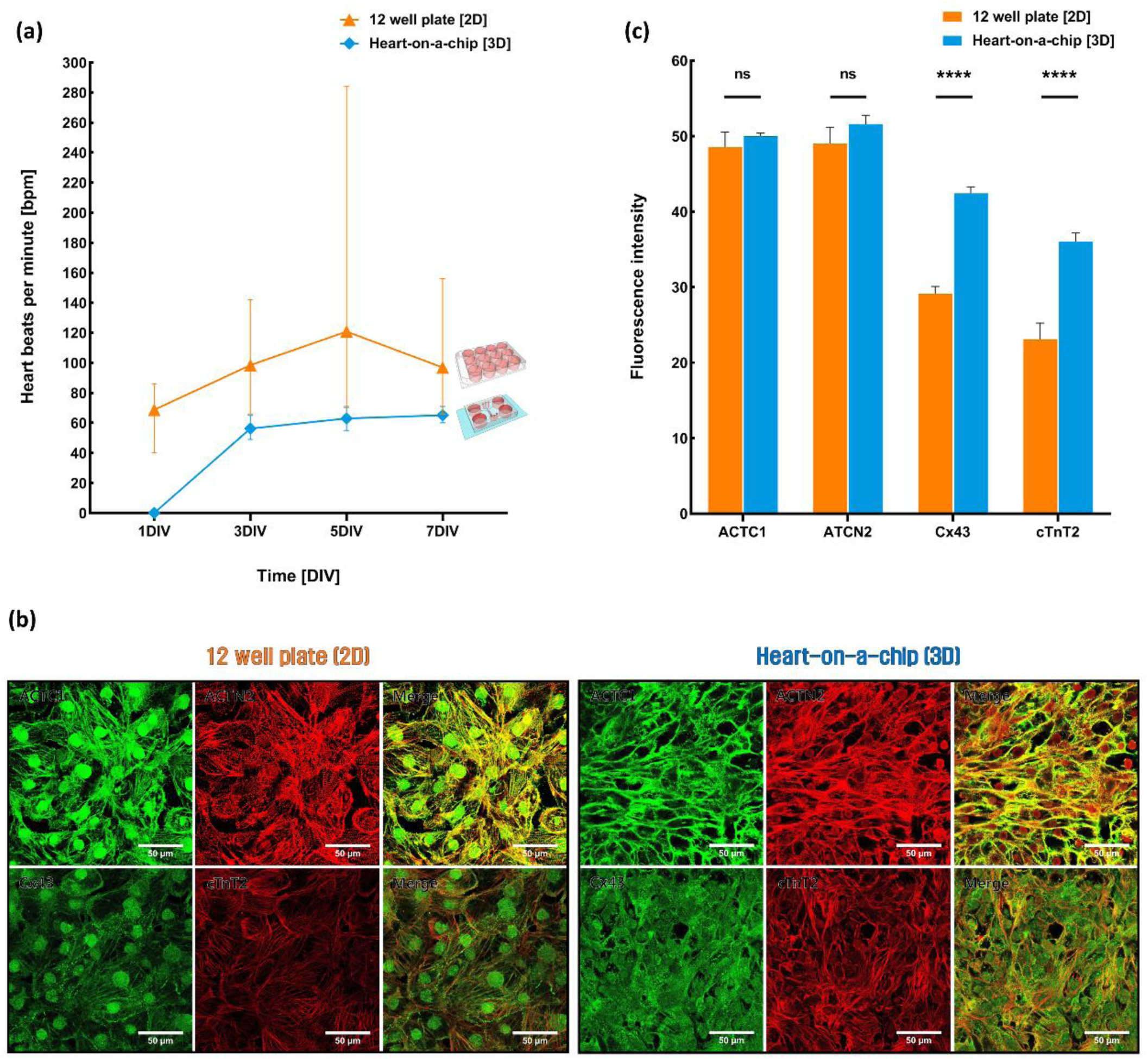
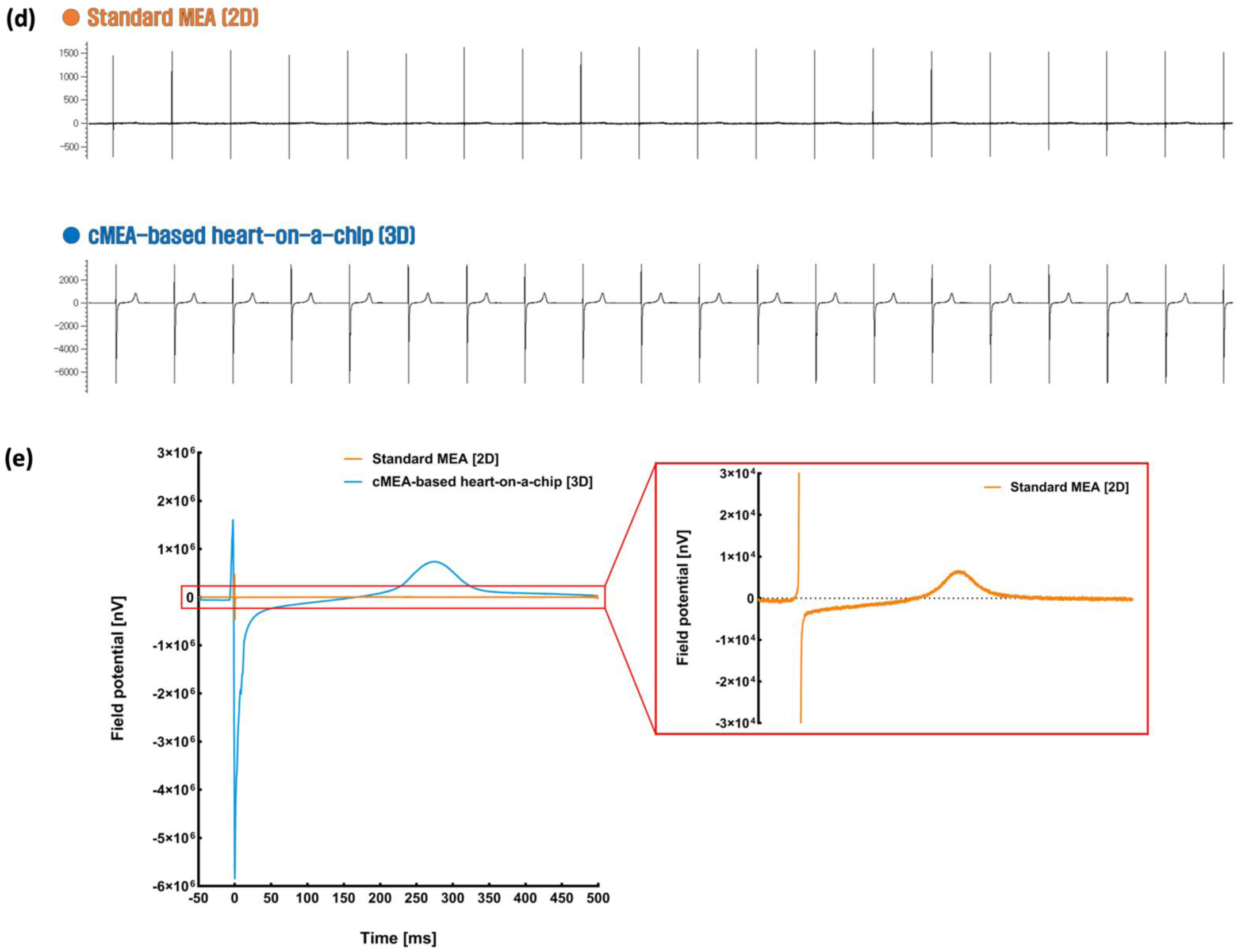
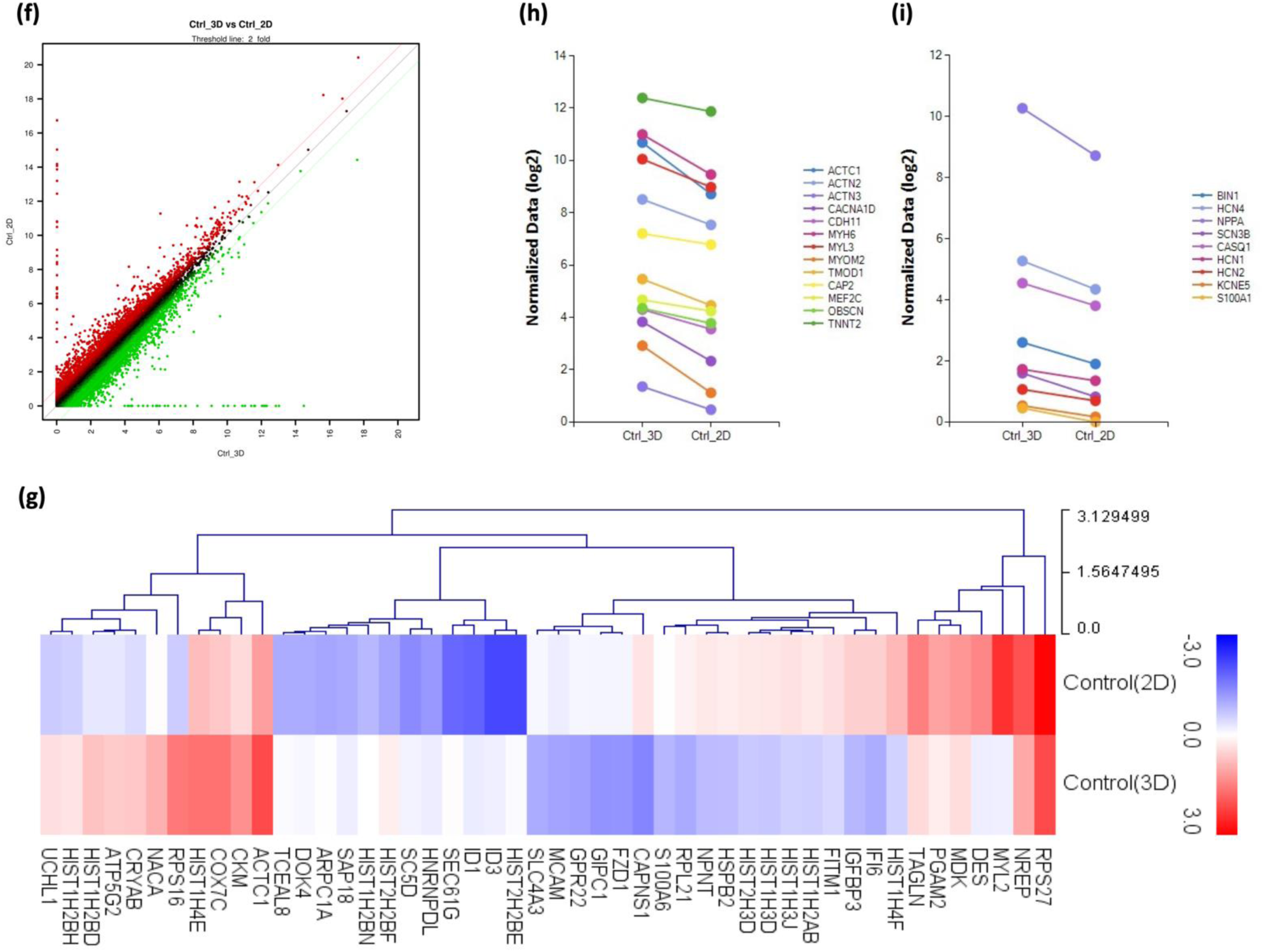
Comparative analysis of hiPSC-CMs on cell culture plate (2D) and heart-on-a-chip (3D). (a) Difference in the heartbeats per minute during culture optimization. (b) Difference in the expression pattern of cardiac markers between the 12 well plate and heart-on-a-chip (cardiomyocyte-specific markers, ACTC1 and Cx43, green; ACTN2 and cTnT2, red). Scale bars: 50 μm. (c) Bar graph of difference in the expression of cardiac markers. Statistical analysis was executed by two-way ANOVA with Tukey’s multiple comparisons test. \*\*\*\**P* <0.0001. Figure 4(a) indicates that the 3D culture of hiPSC-CMs in the chip shows excellent reproducibility. Figures 4(b) and 4(c) indicate that difference in Cx43 expression shows 3D expansion of the contact area between cells by 3D culture. All data are mean ± SD. (d) Electrocardiographic changes of hiPSC-CMs. (e) Scale gap of field potential waveform. Figures 4(d) and 4(e) indicate the contractility at the tissue level with a scale gap of about 100 times compared to the contractile force at the single cell level.(f) Scatter plot graph based on the total RNA-seq data of hiPSC-CMs (log2 mean, p-value < 0.05). (g) Heatmap representing the change for the top 48 differentially regulated genes between 2D and 3D (Fold change > 3, Normalized Data (log2) > 6, p-value < 0.05). (h) Specific gene expression graph related to cardiac structure and function. (i) Specific gene expression graph related to conduction and calcium handling.

#### 2.4.2. Difference in the expression pattern of cardiac markers

We confirmed the expression pattern of crucial markers representing the identity of cardiomyocytes via immunostaining to check the differences in characteristics of hiPSC-CMs cultured in 2D and 3D, along with the preceding comparative analysis of heart rate. Consequently, the expression patterns of ACTC1 and ACTN2 did not show significant changes in 2D or 3D cultures, but noticeable differences of Cx43 and cTnT2 were observed at a glance (**Figure 4b**). When analyzed quantitatively, no significant difference was observed in the fluorescence intensities of ACTC1 and ACTN2 in 2D or 3D culture, but it was confirmed that the fluorescence intensities of Cx43 and cTnT2 were higher in 3D culture compared to 2D culture (**Figure 4c**, ACTC1/2D 48.55; ACTC1/3D 49.98; ACTN2/2D 48.99; ACTN2/3D 51.61; Cx43/2D 29.15; Cx43/3D 42.42; cTnT2/2D 23.1; cTnT2/3D 36.01). However, these quantitative differences in fluorescence signals could not be considered an accurate comparison. Because the nature of 2D culture is that cells are spread widely and attached to the bottom, we should consider that the cell density per same area in 2D culture is lower compared to 3D culture, which may lead to lower fluorescence signals. Instead, the notable point to focus on here was the difference in the expression pattern of Cx43. In 2D culture, the expression was concentrated in the nucleus and the areas where cells come into contact, whereas in 3D culture, the expression was observed throughout the entire cell. This phenomenon can be interpreted as a result of the 3D expansion of the contact area between cells through biomimicking a 3D microenvironment.

#### 2.4.3. Scale gap of field potential waveform

A comparative analysis of electrophysiological aspects according to the 2D and 3D culture of hiPSC-CMs was also performed. We cultured hiPSC-CMs on both conventional MEA, widely used in general, and cMEA integrated into the heart-on-a-chip for 7 days, and then field potential was measured. The results revealed noticeable differences in the scale of the electrocardiogram waveforms (**Figure 4d**). Specifically, when selectively comparing the peak value of the T-wave amplitude in the field potential, 6.57 x 10^3^ nV was measured in the 2D culture, while 7.37 x 10^5^ nV was measured in the 3D culture (**Figure 4e**). These values show a scale gap of approximately 100 times or more, interpreted as the result of the difference in contractility between the single-cell and tissue levels. Therefore, we were convinced that the functionality of the realized myocardial tissue within the heart-on-a-chip has advanced beyond the cellular level, closer to the level of native tissue.

#### 2.4.4. Transcriptomic difference between cell culture plate (2D) and heart-on-a-chip (3D)

The molecular biological changes in hiPSC-CMs according to 2D and 3D cultures were also shown. Differentially expressed genes (DEGs) analysis via total RNA sequencing was performed on a total of 26255 genes in hiPSC-CMs cultured for 7 days. The resulting scatter plot graph in Figure 4f shows the transcriptomic differences between 2D and 3D cultures at a glance. The top 48 protein-related genes that were significantly and differentially regulated between 2D and 3D cultures can be seen in Figure 4g. In detail, compared to 2D cultures, the expression of several genes, including RPS27, NREP, MYL2, DES, MDK, PGAM2, TAGLN, HIST1H4F, IFI6, IGFBP3, FITM1, HIST1H2AB, HIST1H3J, HIST1H3D, HIST2H3D, HSPB2, NPNT, RPL21, S100A6, CAPNS1, FZD1, GIPC1, GPR22, MCAM, and SLC4A3, were decrease. Meanwhile, the expression of several genes, including HIST2H2BE, ID3, ID1, SEC61G, HNRNPDL, SC5D, HIST2H2BF, HIST1H2BN, SAP18, ARPC1A, DOK4, TCEAL8, ACTC1, CKM, COX7C, HIST1H4E, RPS16, NACA, CRYAB, ATP5G2, HIST1H2BD, HIST1H2BH, and UCHL1, were increased.

Within these overall molecular biological changes, we checked references related to transcriptomic analysis on 2D and 3D cultures of cardiomyocytes to determine whether 3D cultures have a positive effect on biomimicking cardiac tissue in particular. Most references suggest that the 3D microenvironment in which cardiomyocytes grow despite each different culture method, such as spheroids, organoids, and biomimetic chips, influences various aspects of cardiomyocyte maturation. Yet, the gene expressions indicating maturity did not show the same pattern [75–81]. Based on our finding, upregulated genes related to maturation in 3D culture were broadly classified as being involved in cardiac tissue structure/function and conduction/calcium handling. Genes involved in processes related to the structure and function of cardiac tissue include ACTC1, ACTN2, ACTN3, CACNA1D, CDH11, MYH6, MYL3, MYOM2, TMOD1, CAP2, MEF2C, OBSCN, and

TNNT2 (**Figure 4h**), and genes involved in processes related to conduction and calcium handling include BIN1, HCN4, NPPA, SCN3B, CASQ1, HCN1, HCN2, KCNE5, and S100A1 (**Figure 4i**). In conclusion, we observed significant changes in the maturity of DEG patterns depending on the culture method. This proves that attempts to realize myocardial tissue through a heart-on-a-chip are meaningful in producing more accurate results.

### 2.5. Drug-induced cardiotoxicity tests of hiPSC-CMs on the heart-on-a-chip

#### 2.5.1. Difference in the drug-induced heart rate

We aimed to confirm whether it was possible to assess the cardiotoxicity of drugs using the heart-on-a-chip. Among the representative drugs that affect heart rate, isoproterenol, which induces faster heartbeats, and nifedipine, which induces slower heartbeats, were administered, and changes in heart rate were intuitively observed. For this heart rate analysis only, we attempted to reaffirm the significance of the heart-on-a-chip in cardiotoxicity evaluation by comparing the drug responses of the 2D cultured cardiomyocytes. After culturing for 7 days, we checked that all chips reached the normal heart rate range of 60-90 bpm and selected some wells within the plate showing heartbeats close to the range, as mentioned above. Isoproterenol was treated at concentrations of 0.1, 1, 5, 10, and 20 μM, and nifedipine was treated at concentrations of 0.1, 1, 5, 20, and 100 nM and then observed at 1 h intervals for a total duration of 24 h.

After Isoproterenol treatment, the heart rate changes in the 12 well plate showed an expected increase in heartbeats at all concentrations 1 h after drug treatment. However, the slight rise in heart rate under the 1 μM condition, despite a concentration higher than 0.1 μM, suggests that dose-dependent changes were not reflected. Besides, the abnormal sudden increases in heart rate, such as reaching 204 bpm, were observed in the 5 μM treatment condition (**Figure 5a**) (**Movie S10**). When examining the changes over time for 24 h, the abnormal fluctuation that the heart rate drastically increased and then decreased regardless of the concentration until 8 h after drug treatment (**Figure S1a**). Considering these results, we inferred that the abnormal reactions to drugs and the difficulty in quantitative analysis make it challenging to perform drug efficacy/toxicity assessments accurately. On the other hand, there was no significant change in the heart rate change in the heart-on-a-chip in the 0.1 μM treatment condition 1 h after drug treatment, but the heartbeats increased under the 1 and 5μM conditions, showing the expected drug efficacy as in the 12 well plate. However, under relatively high concentrations of 10 and 20 μM conditions, the heart rate decreased as the dosage increased, which we interpreted as a result of drug toxicity associated with the high concentrations. Overall, the dose-dependent responses were observed, and no abnormal sudden increases in heart rate occurred at any concentrations (**Figure 5b**) (**Movie S11**). When checking changes for 24 h, except for 10 and 20μM conditions, the heart rate gradually decreased when it reached a certain level after drug treatment. Even in the 10 and 20 μM treatment conditions, the heart rate temporarily stopped or slowed down, and then the heartbeats reached a certain level and decreased as in other concentration conditions. We assumed that the temporary stopping or slowing down phenomena were due to sudden shock caused by high concentration. In summary, the increase or decrease amplitude was larger quantitatively when the concentration was higher, and the heart rate did not rise beyond 110 bpm even under high concentrations (**Figure S1b**). Thus, these results suggested that the 3D culture using the heart-on-a-chip can measure quantitative changes in response to the drug concentrations within the normal heart rate range, compared to the conventional 2D culture, so we believed this provided as evidence that our chip can perform more precise drug efficacy/toxicity evaluation.

**Figure 5.**
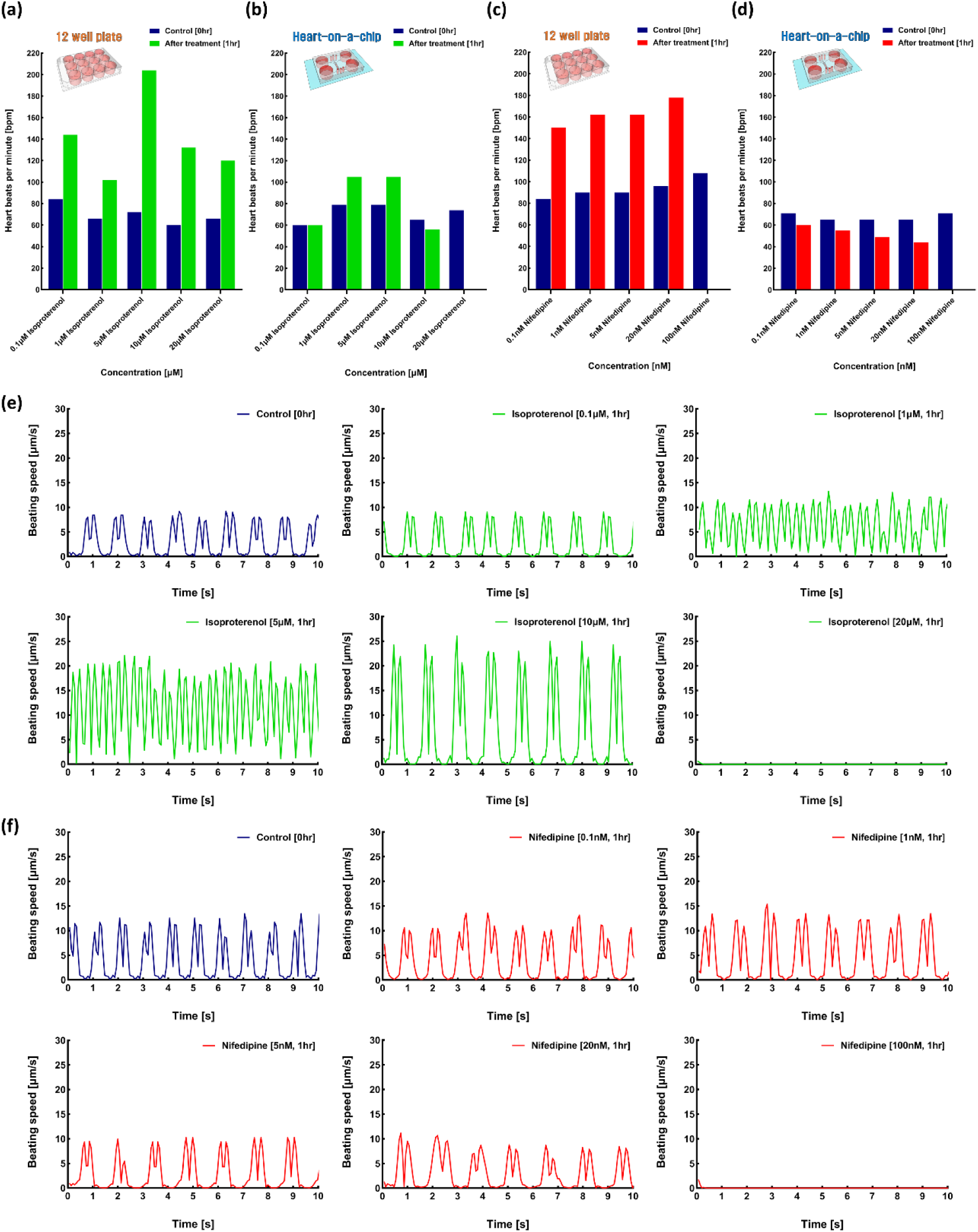
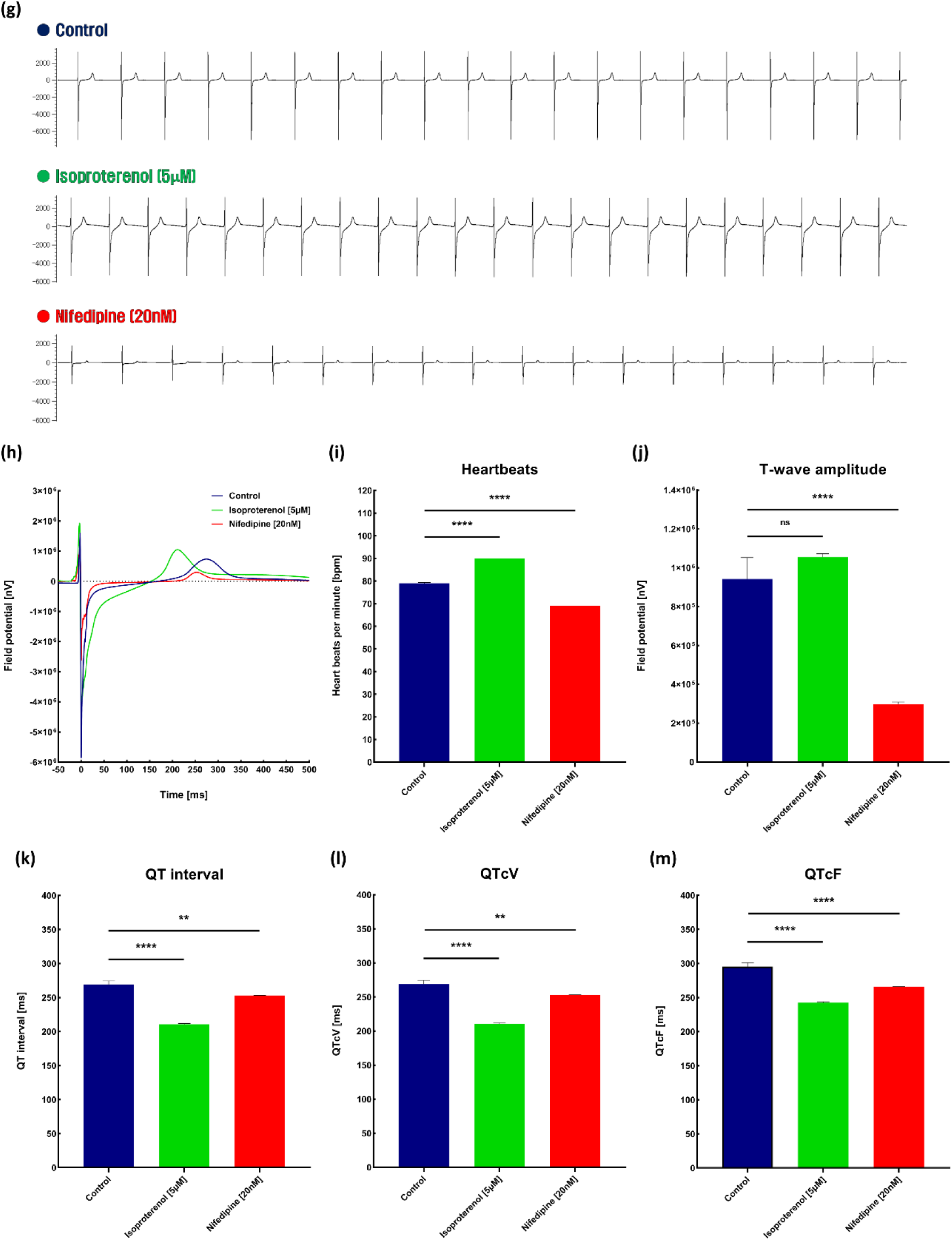
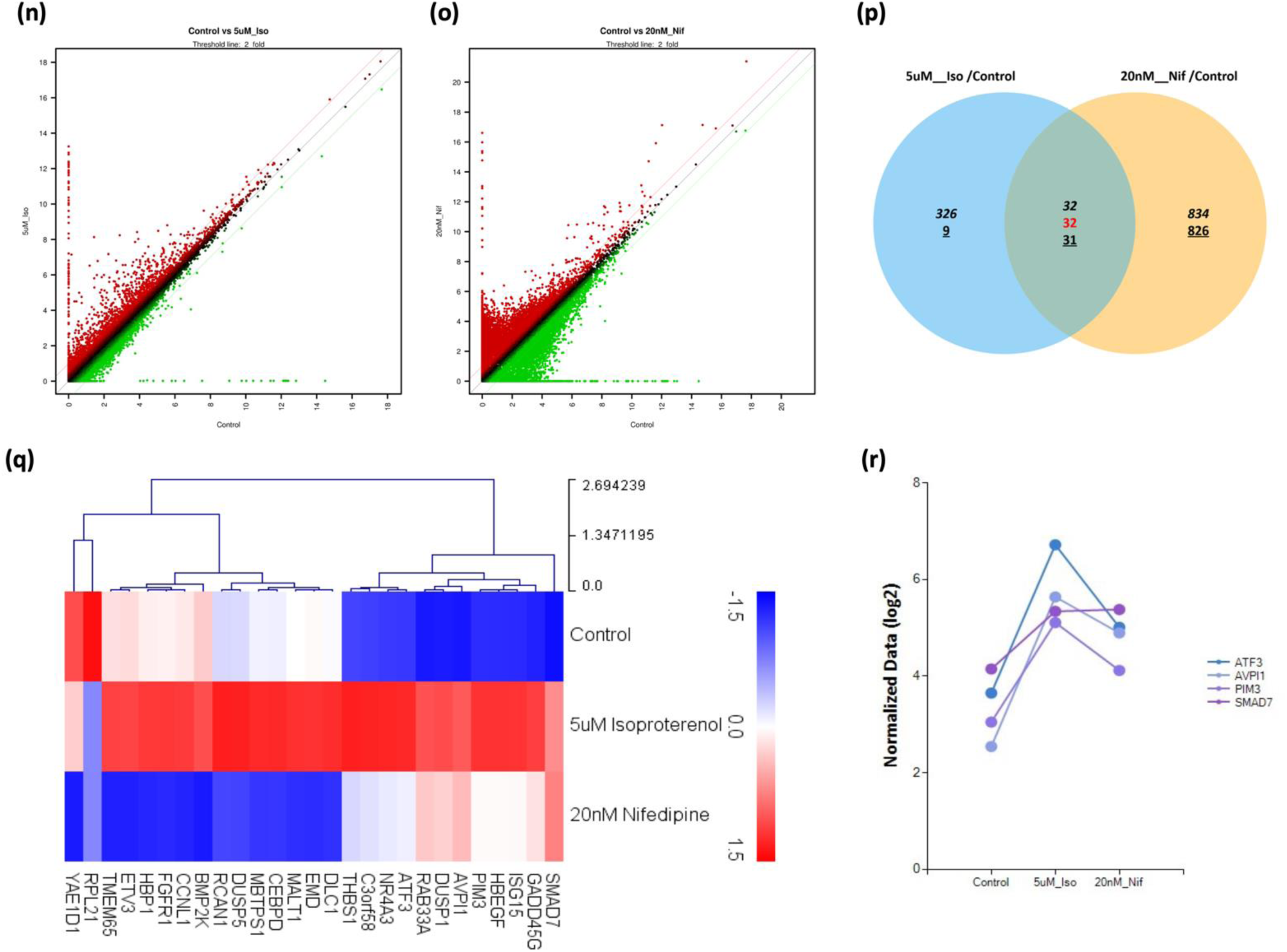
Drug-induced cardiotoxicity tests on the heart-on-a-chip. (a) Heart rate of hiPSC-CMs on the 12 well plate after isoproterenol treatment for 1 h. (b) Heart rate of hiPSC-CMs on the heart-on-a-chip after isoproterenol treatment for 1 h. (c) Heart rate of hiPSC-CMs on the 12 well plate after nifedipine treatment for 1 h. (d) Heart rate of hiPSC-CMs on the heart-on-a-chip after nifedipine treatment for 1 h. (e) Motion tracking of hiPSC-CMs on the heart-on-a-chip after isoproterenol treatment for 1h. (f) Motion tracking of hiPSC-CMs on the heart-on-a-chip after nifedipine treatment for 1 h. (g) Effects of drugs on Electrocardiogram of hiPSC-CMs. (h) Effects of drugs on Field potential of hiPSC-CMs. (i) Effects of drugs on MEA-based heart beats per minute of hiPSC-CMs. (j) Effects of drugs on T-wave amplitude of hiPSC-CMs. (k) Effects of drugs on QT interval of hiPSC-CMs. (l) Effects of drugs on QTcV of hiPSC-CMs. (m) Effects of drugs on QTcF of hiPSC-CMs. Statistical analysis were done using one-way ANOVA with Tukey’s multiple comparisons test. \*\**P* = 0.0067; \*\*\*\**P* <0.0001. All data are mean ± SD. (n) Scatter plot of the differentially regulated genes between control and isoproterenol treatment (log2 mean, p-value < 0.05) (o) Scatter plot of the differentially regulated genes between control and nifedipine treatment (log2 mean, p-value < 0.05). (p) Heatmap representing the change for the top 27 differentially regulated genes depends on drug treatments (Fold change > 2, Normalized Data (log2) > 4, p-value < 0.05). (q) Venn diagram based on the RNA-seq data depends on drug treatments (Fold change > 2, Normalized Data (log2) > 4, p-value < 0.05). (r) Selected gene plot graph based on the total RNA-seq data depends on drug treatments.

Meanwhile, in the case of nifedipine, contrary to the expected effect of slowing down heartbeats at all concentrations, the heart rate changes in the 12 well plate abnormally rose to over 150 bpm except for the 100 nM condition after 1 h of drug treatment. The heartbeats temporarily stopped in the highest concentrated condition of 100 nM. This temporary pause was also assumed to be caused by sudden shock because of high concentration. Although it appeared contrary to the expected drug efficacy, quantitative changes depending on the concentration were observed, unlike when treated with isoproterenol (**Figure 5c**) (**Movie S12**). When observing changes over 24 h, heartbeats drastically increased and gradually decreased when they reached a certain level. Only the 100 nM treatment showed recovery from the sudden shock-induced temporary arrest in 3 h and thereafter exhibited a decrease similar to other concentrations (**Figure S1c**). The results, contrary to the well-known drug effects and the heart rate far exceeding the normal range, had once again confirmed the difficulty of accurately conducting drug efficacy/toxicity assessments through the conventional culture methods. On the other hand, the changing heart rate pattern in the heart-on-a-chip accurately represented the drug efficacy by exhibiting a dose-dependent decrease in heart rate for all concentration conditions after 1 h of drug treatment. Similar to the 12 well plates, the temporary stoppage of heartbeats owing to sudden shock was observed only under 100 nM conditions (**Figure 5d**) (**Movie S13**). When checking the changing pattern with the lapse of time up to 24 h, the heart rate, which was temporarily stopped or slowed down in all concentration conditions, was restored to a normal level when 6 h had elapsed after drug administration. Exceptionally, in the 100 nM treatment condition where the heart rate temporarily disappeared, the intensity or speed of the heartbeats did not return to normal levels, although the heartbeats recovered (data not provided). As time elapsed close to 24 h, a gradual decrease in heart rate was also observed as the drug was treated at a high concentration (**Figure S1d**). Therefore, similar to isoproterenol, these results suggest that the 3D culture in the heart-on-a-chip is expected to facilitate more accurate drug efficacy/toxicity evaluations in all aspects compared to traditional 2D culture.

#### 2.5.2. Drug-induced beating pattern changes with motion tracking analysis

The changes in heart rate due to the drugs can be visually confirmed through simple counting, but other changes in heartbeat patterns, such as the speed or intensity of each heartbeat, are difficult to estimate through intuitive observation alone. Hence, we attempted to prove that various results can be obtained through video-based motion tracking based on the heart-on-a-chip to check not only the heart rate but also the changes in the heartbeat pattern after drug treatment. For this purpose, we selected the previous video results of heart rate changes observed after 1h treating isoproterenol at concentrations of 0.1, 1, 5, 10, and 20 μM and nifedipine at concentrations of 0.1, 1, 5, 20, and 100 nM. The selected videos were analyzed through motion tracking, and variations in heart rate according to the type and concentration of the drugs were graphed.

The heartbeat pattern for isoproterenol treatment in control without drug treatment exhibited regular contractions/relaxations with an average speed of approximately 7.5 μm/s, maintaining a consistent intensity and interval. After drug treatment, it responded dose-dependently. There was no significant change under the 0.1 μM concentration, while at 1 and 5 μM, the heartbeat speed increased to approximately 10 and 18.5 μm/s, respectively. The pulse interval also narrowed proportionally to the administered amount of the drug. In the previous heart rate analysis, a concentration of 10 μM led to a decrease in heart rate due to shock. In the motion tracking analysis, the consistent pattern that the heartbeats slowed as the interval increased was confirmed, but the contractions’ amplitude increased even further, reaching an average speed of about 23 μm/s. These results indicate the potential for deriving more intricate results that cannot be obtained through solely manual heartbeat counting, as showing the heartbeat intensity became stronger, attributed to the inherent efficacy of the drug, despite a decrease in heart rate due to shock from the higher concentration. In the remaining 20μM concentration condition, no waveform was created because of the disappearance of the pulsations (**Figure 5e**) (**Movie S11**).

The beating pattern in response to nifedipine in control without drug treatment showed maintained contractions/relaxations with a regular amplitude and interval at an average speed of about 10 μm/s. After drug treatment, although the reactions were subtler than the responses with isoproterenol, the quantitative reactions were shown. At concentrations of 0.1 and 1 nM, there was no significant change in the heartbeat intensity, but the interval between pulsations slightly increased. Under 5 nM and 20 nM, the heartbeat speed decreased to an average of approximately 8.5 and 7 μm/s, respectively, and the interval widened. No waveform was formed in the remaining 100 nM treatment condition because the heartbeats disappeared (**Figure 5f**) (**Movie S13**). Consequently, this cardiac biomimetic chip demonstrated the potential as a more precise evaluation tool for cardiotoxicity by allowing motion tracking analysis, thereby enabling the observation of detailed drug responses.

#### 2.5.3. Drug-induced electrophysiological alterations

In the case of the heart that exhibits electrophysiological characteristics like action potential, functional assessment is essential. Hence, we aimed to evaluate whether electrophysiological changes caused by drugs could also be detected, as well as the previous video-based analyses. First, hiPSC-CMs were cultured for 7 days on the heart-on-a-chip integrated with cMEA. Subsequently, we chose 5 μM isoproterenol and 20 nM nifedipine treatment conditions, which showed the most noticeable changes in dose-dependent experiments, and then the field potential was measured 1 h after drug treatment at selected concentrations. As a result, alterations in electrocardiogram waveforms were clearly observed depending on the type of drug (**Figure 5g and 5h**). Furthermore, based on these measurements, we conducted a more detailed analysis using three evaluation parameters: heart rate, T-wave amplitude, and QT interval. In particular, the QT interval is one of the rating indices of the electrocardiogram used to assess the electrophysiological characteristics of the heart and means the time from the start of the QRS complex to the end of the T wave. This also refers to the depolarization and repolarization period of the ventricles, that is, the time taken from the onset of contraction to the completion of relaxation. Since an abnormally long or short QT interval can cause abnormal heart rhythm and sudden cardiac death [82–86], it is a critical evaluation criterion for cardiotoxicity. Because this QT interval depends on changes in heart rate, it was converted for values not influenced by heart rate changes using QT interval correction formulas such as Van der Linde’s (QTcV) and Fridericia’s (QTcF). The heart rate increased after treatment with 5 μM isoproterenol and decreased after treatment with 20 nM nifedipine, compared to the control group (**Figure 5i**, control 79 bpm; 5 μM isoproterenol 90 bpm; 20 nM nifedipine 69 bpm). These results demonstrated reproducibility regardless of the analytical methods, as they were consistent with the video-based analysis of beating pattern changes, such as manual heartbeat counting and motion tracking performed previously. T-waveform amplitude, compared to the control, slightly increased when treated with 5 μM isoproterenol and significantly decreased when treated with 20 nM nifedipine (**Figure 5h and 5j**, control 9.42 x 10^5^ nV; 5 μM isoproterenol 10.54 x 10^5^ nV; 20 nM nifedipine 2.97 x 10^5^ nV). These results also showed that changes in the contraction/relaxation force induced by the drugs could be observed, similar to the beating pattern analysis through motion tracking. Finally, the QT interval, QTcV, and QTcF were confirmed to be shorter in both conditions treated with 5 μM isoproterenol and 20 nM nifedipine compared to the control (**Figures 5h, 5k, 5l, and 5m**, control/QT interval 268.82 ms; 5 μM Isoproterenol/QT interval 210.78 ms; 20 nM nifedipine/QT interval 252.9 ms; control/QTcV 268.84 ms; 5 μM Isoproterenol/QTcV 210.79 ms; 20 nM nifedipine/QTcV 252.91 ms; control/QTcF 295.47 ms; 5 μM Isoproterenol/QTcF 242.55 ms; 20 nM nifedipine/QTcF 265.85 ms). These results accurately reflect nifedipine characteristics that do not lengthen the QT interval even as the heart rate decreases as a non-QT prolonging drug, unlike isoproterenol, in which the QT interval naturally shortens as the heart rate increases. In conclusion, we have proven that the application of MEA technology to the heart-on-a-chip can precisely analyze electrophysiological evaluation parameters, such as QT interval, which cannot be confirmed through video-based analysis alone. Hence, The superior functionality of the heart-on-a-chip shows that it is a platform that can adequately meet the purpose of new pharmacological safety evaluations such as CiPA.

#### 2.5.4. Transcriptomic difference after isoproterenol and nifedipine treatments

We also observed molecular biological changes in hiPSC-CMs upon treatment with different types of drugs. After drug treatment, hiPSC-CMs were recovered from the chips, and DEG analysis was carried out by total RNA sequencing for 26255 genes as in the previous analysis. The scatter plot graphs in Figures 5n and 5o show the overall transcriptome alterations following isoproterenol and nifedipine treatment compared to the control. Venn diagram based on gene lists with 2fold, normalized data (log2) > 4, and p-value < 0.05 indicates that 324 genes are upregulated and 9 genes are downregulated by isoproterenol treatment, while 834 genes are upregulated and 826 genes are downregulated by nifedipine treatment. We can also see that 32 genes are regulated by both drugs (**Figure 5p**). The top 27 protein-related genes that were notably and differentially regulated by the treated drugs can be seen in Figure 5q. In detail, all genes were significantly upregulated under isoproterenol treatment conditions, albeit to varying degrees, except for RPL21 and YAE1D1, which were downregulated compared to the control group. In contrast, in nifedipine-treated conditions, the expression of genes, including SMAD7, GADD45G, ISG15, HBEGF, PIM3, AVPI1, DUSP1, RAB33A, ATF3, NR4A3, C3orf58, and THBS1 was upregulated compared to the control, but was relatively less upregulated than under isoproterenol treatment conditions. In addition, the expression of genes including DLC1, EMD, MALT1, CEBPD, MBTPS1, DUSP5, RCAN1, BMP2K, CCNL1, FGFR1, HBP1, ETV3, and TMEM65 was significantly downregulated. A direct comparison of the changes for each drug on the graph has shown a clearer difference.

Among the genes that are significantly regulated by drug treatment, the widely studied is activating trans factor 3 (ATF3), an immediate early gene (IEG) that responds to cellular stress. Isoproterenol treatment to cardiomyocytes is known to increase ATF3 expression, and ATF3 overexpression is associated with cardiac hypertrophy. There are also a number of cases in which treatment with large amounts of isoproterenol induces cardiac hypertrophy [87–100]. Nifedipine has also been implicated in cellular stress and is therefore assumed to increase ATF3 expression. Moreover, isoproterenol increases PIM3, AVPI1, and SMAD7 expression, and overexpression has been shown to induce hypertrophy, heart failure, or myocardial ischemia [101–105]. The same results as these basic studies can be seen in Figure 5r. In conclusion, we observed significant changes in DEG patterns depending on the type of treated drugs. While we cannot interpret all of these transcriptomic changes at this time, we believe they are worthy of further basic research. These results also suggest that drug evaluation can be performed more accurately using our heart-on-a-chip.

## 3. Discussion

Cardiotoxicity arising from drugs has a risk of leading to sudden death as the heart is an organ directly related to life, so it must be examined from the drug developmental stage. Particularly considering that such issues occur in many cases when used as medicines for not only heart diseases but also other diseases, cardiac toxicity evaluation is essential for most drugs. Yet, each conventional 2D cell culture and animal test used for this purpose have poor reproducibility and clear limitations of inter-species differences, so drug evaluation methods using them have a high failure rate despite enormous cost and time. Therefore, there was a need for a novel platform that could accurately predict cardiophysiological changes caused by drugs. For these reasons, we developed the hiPSC-CMs-based heart-on-a-chip integrated with the cMEA, which can exhibit excellent reproducibility and comprehensively observe drug-induced cardiotoxic reactions. However, there is still room for complements to derive more meaningful results through the chip developed in this study. As part of the follow-up study, we should execute other experimental conditions that are not verified and additionally apply important biomimetic points that are not reflected. Although we have demonstrated the possibility of cardiac toxicity evaluations with isoproterenol and nifedipine using the chip, it is necessary to appraise the chip with more drugs. For example, if 28 drugs classified into three categories by risk levels in the recently spotlighted CiPA project are assessed using the chip [106], the reliability would be further enhanced as a cardiotoxicity test platform. There is a limit to the maturity of cardiomyocytes that can be reached from hiPSCs through general differentiation methods. In judging CM maturation, there are significant parameters such as elongated rod cell shape, directional alignment of myofibrils, and sarcomere M-line formation [107–109]. Unfortunately, this level of maturity was not attained despite 3D culture using the chip. Hence, in follow-up studies, it is necessary to apply additional conditions, such as physical or electrical stimulation, known to improve CM maturation. It is also necessary to actualize vascularized myocardial tissue via co-culture with endothelial cells and fibroblasts, which are the surrounding cells of cardiomyocytes in vivo. To this end, a sufficient amount of endothelial cells should be loaded into the space between the micro-posts via media channels within the chip. Subsequently, fibroblasts secreting cytokines and ECM should be injected into the chamber with hiPSC-CMs to induce angiogenesis from the media channels toward the chamber, thereby recapitulating myocardial tissue formed with a capillary bed. Furthermore, achieving excellent reproducibility of angiogenesis within the chip requires detailed adjustments, such as optimizing the media composition for co-culture and ideal infusing conditions for tissue vascularization. Lastly, it requires further expanding the roles of the perimysium-like collagen layer inside the chip. Currently, the in-chip collagen layers are limited to minimizing shear stress by inducing media flow similar to diffusion, but more related physiological phenomena can be observed by enhancing chip design elements. For instance, the perimysium is also known to carry out roles such as transmitting lateral contractile force and preventing adhesion between the cardiac fascicles in vivo [50], so if the collagen layers placed in the intervals of plural cardiac fascicles are actualized in the chip, an improved platform that represents more specific physiological characteristics could be established. Furthermore, it is known that cardio-pathological changes, such as thickening of the perimysium, occur depending on cardiac disorders [110]. Therefore, it will be possible to utilize it for heart disease research via chip-based disease modeling and analysis of the collagen layer components. As such, we hope that the heart-on-a-chip will be used as a complete alternative to animal experiments in cardiotoxicity tests by reaching a certain level that can perfectly recapitulate the physiological characteristics of the heart that cannot be realized with current technology over time.

## 4. Conclusion

We developed the hiPSC-CMs-based heart-on-a-chip integrated with the cMEA and confirmed the significance of 3D tissue culture compared to the traditional 2D cell culture. We also validated whether or not cardiotoxicity evaluation using the chips is possible in multilateral means such as kinetic, electrophysiological, and molecular biological analyses. In detail, the asymmetrically rounded rectangular-shaped micro-posts with a zipper-like zigzag arrangement allowed viscous fluids to be separately injected into adjacent spaces without mixing. Applying the silicone adhesive to the fabrication of microfluidic units made reversible bonding with simple attachment and neat detachment practicable, enabling the harvest of cardiac tissues formed in the chip and the reuse of the cMEA electrode units. The MEA electrode units were customized to be compatible with the heart-on-a-chip for electrophysiological measurements. The myocardial fascicle, including endomysium and perimysium, and the surrounding capillaries were three-dimensionally actualized in the chip using human genome-based iPSC-CMs and collagen. It was checked why the 3D-cultured within the chip is necessary compared to the 2D-cultured in the plate through the constant heart rates, the difference in Cx43 expression pattern, the field potential waveform scaled 100 times, and the gap of the DEGs. Isoproterenol and nifedipine were treated as representative drugs capable of controlling the heart rate, and then the efficacy/toxicity tests were carried out dose-dependently as time advanced. Afterward, by observing the change in the heart rate, we affirmed that the 3D culture using the chip is available to present more accurate results than the traditional 2D culture. Furthermore, we analyzed kinetic results like the speed (or intensity) and pattern of each heartbeat through video-based motion tracking, specific electrophysiological results including the electrocardiogram waveform, heart rate, T-wave amplitude, R-R waves interval mean, and QT interval by field potential measure, and molecular biological results such as the DEG patterns altered by the drugs via transcriptome profiling. Through this, we proved that the chip could be utilized to analyze various indicators that cannot be evaluated only by intuitive heart rate analysis. Conclusionally, we expect our heart-on-a-chip will become a well-timed cardiotoxicity assessment platform based on good reproducibility and multifaceted analysis at a time when new pharmacological safety evaluations, such as CiPA, have been gaining attention recently.

## 5. Experimental Section

### Fabrication of cMEA

The cMEA was designed through an AutoCAD program (Autodesk) and fabricated through a standard photolithography process **(Figure S2)**. A transparent chrome mask for electrode patterning was first fabricated on a 5-inch quartz glass substrate using an E-beam lithography system (JEOL, JBX-6000 FS). Then, titanium (Ti), constituting a conductor such as tracks, was deposited using an E-beam evaporator (ULTEC, UEE) on a glass substrate, and titanium nitride (TiN), constituting electrodes and contact pads, was deposited using a cluster sputter (ULVAC, SME-200J). The blank glass substrate with the thin metal film was loaded with 1.5 ml of positive photoresist (PR) (AZ Electronic Materials, AZ7220 (33CP)) and spin-coated using a spin process controller (MIDAS, Spin-3000D). The positive PR-coated glass substrate was soft-baked on a 110 °C hot plate for 90 sec. After soft baking, the positive PR-coated glass substrate was covered with the pre-fabricated chrome mask and exposed to UV using a mask aligner (MIDAS, MDA-400M), and after UV exposure, post-exposure baking (PEB) was performed on a 110 °C hot plate for an additional 90 sec. The glass substrate with the patterned electrode was immersed in AZ300 MIF developer (AZ Electronic Materials) to develop UV-exposed PR for 90 sec, and immersed in TiN 22-7 titanium nitride etchant (Transene) for 60 sec to etch TiN, followed by immersion in TFTN titanium etchant (Transene) for 90 sec to etch Ti. Finally, the remaining PR on the glass substrate was removed with acetone. Each etching process step was rinsed with deionized water (DI water) and dried using a nitrogen gun. Lastly, silicon nitride (SiN) was coated as an insulator on all parts except the electrode and contact pad using a low-density plasma-enhanced chemical vapor deposition (PECVD) (Unaxis, VL-LA-PECVD) equipment.

### Fabrication of microfluidic units

The microfluidic unit was also designed through the AutoCAD program (Autodesk) and fabricated using PDMS as the base material through a soft lithography process (**Figure S3**). Briefly, a transparent chrome mask for microstructure patterning of microfluidic units was first fabricated using an E-beam lithography system (JEOL, JBX-6000 FS) on a 5-inch quartz glass substrate. After that, negative PR (MicroChem Corp, SU-8) was spin-coated on a silicon wafer with a thickness of 50 μm for mold fabrication and soft-baked on an 80 °C hot plate for 10 min. After soft baking, the negative PR-coated silicon wafer was covered with the chrome mask prepared in advance and exposed to UV using a mask aligner (MIDAS, MDA-400M). The UV-exposed silicon wafer was immersed in SU-8 developer (Micro Chem Corp), developed for 90 sec, rinsed with isopropyl alcohol, and dried with nitrogen. The patterned wafer was post-baked (PEB) on a hot plate at 80 °C for 10 min. PDMS (DuPont, Sylgard 184) was mixed with a curing agent in a ratio of 10:1, poured into a mold patterned with negative PR, and cured on a hot plate at 80 °C for 24 h. The cured PDMS-based microfluidic units were peeled off from the mold and checked for microstructural abnormalities. Inlets and outlets for cell and collagen solution injection were created using a 1.5 mm biopsy puncher (Ted Pella), and wells for media reservoirs were created using an 8 mm biopsy puncher (World Precision Instruments). Debris attached to the PDMS-based microfluidic units was simply removed with tape, soaked in isopropyl alcohol, washed with ultrasonication for 60 sec, and then dried completely in a dry oven at 80 °C for 10 min. For sterilization for cell culture, the completed microfluidic units were exposed to UV light for 1 h.

### Silicone adhesive application for reversible bonding of microfluidic units

For reversible bonding, silicone adhesive was applied to PDMS to provide adhesion to the microfluidic units themselves, thereby replacing irreversible bonding such as oxygen plasma bonding. First, the PDMS and curing agent were mixed and prepared in an 8:1 ratio to a volume of 12.5 g. The silicone adhesive components, Parts A and B, were mixed and prepared in a 1:1 ratio to a volume of 37.5 g. Finally, the PDMS:silicone adhesive mixture was mixed and prepared in a 1:3 ratio to a volume of 50 g. After removing air bubbles generated during mixing under vacuum, this mixture was poured into a mold patterned with negative PR coated with 5% pluronic F127 (Sigma) and cured on a hot plate at 80 °C for 24 h. The cured PDMS:silicone adhesive-based microfluidic units were peeled off from the mold, and the inlets, outlets, and media reservoirs were created using the 1.5 mm biopsy puncher and the 8 mm biopsy puncher. The PDMS:silicone adhesive-based microfluidic units were cleaned by simply removing debris with tape, immersed in isopropyl alcohol, sonicated for 60 sec, and then completely dried in a dry oven at 80 °C for 10 min. To sterilize for cell culture, the completed microfluidic units were exposed to UV light for 1 h.

### Chip assembly

Depending on the purpose of the experiment, a PDMS or PDMS:silicone adhesive-based microfluidic unit was selected, and the chip was assembled by plasma bonding or thermal bonding by matching one of the substrates: slide glass, cover glass, or cMEA (**Figure S3**). As with the microfluidic units, slide glasses, cover glasses, and cMEAs were first sterilized by simply removing debris with tape, immersing them in isopropyl alcohol, washing them with ultrasonication for 60 sec, drying them, and exposing them to UV for 1 h. Then, the PDMS-based microfluidic units and slide glasses were selected for chips for general experiments, plasma-treated at 40 W for 40 sec using an oxygen plasma treatment system (Femto Science, CUTE-1MPR), and then irreversible bonding was performed on an appropriate position on the substrate. On the other hand, the PDMS:silicone adhesive-based microfluidic units and cMEAs or cover glasses were selected for electrophysiological measurement, immunostaining, and cell harvest, and reversible bonding was performed on an appropriate position on the substrate by heating at 120 °C on a hot plate for 24 h. After plasma and thermal bonding, the chips were finally placed in a dry oven at 80 °C to ensure the reproducibility of separation loading.

### Confirmation of fluid mixing in the microfluidic chip

The feasibility of separate injections into the chip without microvalves and without fluid mixing was confirmed by injecting food coloring into each microstructure with a syringe pump (Harvard Apparatus, Pump 11 Elite). Food coloring of different colors to be injected was prepared by diluting the stock solution 10-fold in sterile water. Only the food coloring to be injected into the perimysium-like collagen layer channel was diluted in collagen solution (Advanced BioMatrix) in the same ratio, considering the physical properties of the actual fluid to be injected. The prepared food coloring solutions were injected through the syringe pump at a rate of 3 μl/min for the perimysium-like collagen layer channels, 20 μl/min for the cardiomyocyte and endomysium chambers, and 10 μl/min for the capillary-like media channels. Whether the food coloring was infused into the desired microstructure without mixing was checked using an inverted microscope (Olympus, CKX41) equipped with a digital camera for the microscope (Olympus, EP50).

### Electrochemical characterization of cMEA

To ensure that the cMEA, which was customized for compatibility with the heart-on-a-chip, functions properly, the sensitivity of the electrode was evaluated by measuring the impedance change according to the presence or absence of hiPSC-CMs. hiPSC-CMs were cultured for 7 days on the cMEA which was arranged in the form of a 60-electrode MEA with all electrodes located in the cardiomyocyte and endomysium chamber of the heart-on-a-chip. The impedance changes in the presence and absence of cells were detected using an in vitro recording system (Multichannel systems, MEA2100-system), and the results were measured and analyzed using the MEA-IT program (Multichannel systems).

### Culture of hiPSC-CMs

Cardiosight-S^®^ (NEXEL), hiPSC-CMs, were purchased, and a protocol was established to ideally seed and culture them on the heart-on-a-chip (**Figure 3a**). Briefly, VitroCol^®^ (Advanced BioMatrix), a human type I collagen, was mixed with 10-fold concentrated RMPI 1640 medium (Gibco), and then it was neutralized with 0.1 M NaOH solution (Millipore) to prepare the collagen solution at a concentration of approximately 3 mg/ml according to the manufacturer’s instructions. PhotoCol^®^ (Advanced BioMatrix), which is type I methacrylated collagen referred to as ColMA for short, was also dissolved in 20 mM acetic acid and neutralized to pH with a neutralizing solution to prepare the ColMA solution at a concentration of about 8 mg/ml. The acetic acid and neutralizing solution are kit components. Each hydrogel solution was prepared at low temperature on ice for at least 1 h before injection into the chip, thereby ensuring an optimal state for 3D gelation. In general, the human-derived collagen solution is mainly used, but when collagen layers with a stronger hardness were desired only for the on-chip collagen layer formation, the ColMA solution was also utilized. The collagen or ColMA solution prepared at low temperature was injected at a low speed of 3 μl/min using the syringe pump through the inlets connected to two perimysium-like collagen layer channels. The injected chips were placed at 37 °C for about 2 h to perform temperature-based gelation primarily. In the case of injection of ColMA, UV-based gelation could further proceed, but we did not carry it out because it showed sufficient hardness for the experiment only with a relatively higher concentration than the existing human-derived collagen solution. Next, a cell-collagen mixture was prepared by mixing 2 x 10^6^ of hiPSC-CMs with 2 μl of human-derived collagen solution and loaded 2 μl of the mixture which included 3.2 x 10^5^ of hiPSC-CMs per each chip at the inlet connected to the cardiomyocyte and endomysium chamber using a pipette. After that, a narrow end of a 1000 μl tip was inserted tightly into the inlet and pushed using the pipette, or tubing was connected to the inlet and pushed through at a rate of 20 μl/min using the syringe pump. If the density of injected hiPSC-CMs was low, optionally, tubing was connected to the outlet connected to the cardiomyocyte and endomysium chamber and sucked up at a rate of 100 μl/min using the syringe pump. The cell density in the chamber was ideally controlled in such a way that the cells were loaded in a micro-weir structure and the collagen solution was selectively removed under the micro-weir. The infused chips were placed at 37 °C for about 30 min to perform secondary temperature-based gelation. Thereafter, each 200 μl of Cardiosight-S^®^ media (NEXEL) that were added Cardiosight-S^®^ supplement (NEXEL) diluted 100 times was first loaded on the two inlet wells among a total of four wells that are media reservoirs linked with two capillary-like media channels and pushed using the pipette toward the remaining empty outlet wells connected to the channels, followed by loading additional 200 μl of the media to ensure a smooth flow of media between the channels and the media reservoirs. In all injection processes, the temporarily formed gaps between the chamber and the channels were removed by allowing them to sit at room temperature for about 30 min, thereby ensuring the media flowed through all microstructures in the chip. The chips prepared in this way were cultured in the same media for 7 days at 37 °C in a humidified 5 % CO^2^ incubator, with the media replacement method changed according to how the heart rate changed with the incubation time. Since the heartbeat of the hiPSC-CMs in the chip began around day 2 of culture, all media reservoirs were loaded with 200 μl of media for the first 2 days of culture to minimize media flow into the chip microstructure and allow the cells time to adapt to the new on-chip environment. From the time the heartbeats were observed, 200 μl of the media was loaded into the two inlet wells, and 100 μl of the media was loaded into the two outlet wells to give a difference in the height of the media to induce gravitational media flow. The media was changed every 2 days and the optimal incubation period was set at 7 days to reach the normal heart rate range of 60-90 bpm based on heart rate. For comparative analysis with cardiomyocytes in conventional 2D culture, Cell-culture treated 12 well plate (NUNC) was additionally coated with Matrigel® hESC-Qualified (Corning) and 3.2 x 10^5^ of hiPSC-CMs per well were loaded using the pipette and cultured for 7 days under the same conditions.

### Immunocytochemistry

Immunocytochemical staining was performed via the following antibodies to check the characteristics of cardiomyocytes cultured in 3D on the heart-on-a-chip and at the same time to compare with cardiomyocytes cultured in 2D: anti-actin cardiac muscle 1 (ACTC1) (1:50) (GeneTex), anti-actinin alpha 2 (ACTN2) (1:1000) (Sigma), anti-connexin 43 or gap junction protein alpha 1 (Cx43 or GJA1) (1:200) (GeneTex), anti-cardiac troponin T or Troponin T2 cardiac type (cTnT2 or TNNT2) (1:250) (GeneTex). Briefly, hiPSC-CMs cultured on the heart-on-a-chips and 12 well plates, respectively, were fixed with 10 % formalin solution (Sigma) for 1 h. After that, in the case of the chip, the microfluidic units interfered with immunostaining, so they were removed from the substrates and wells were temporarily created for fluid loading. The fixed cells to the chips and plates were permeabilized in 0.5 % Triton X-100 (Sigma) for 48 h. A blocking buffer was prepared by mixing 1 % bovine albumin serum (BSA) (Gibco) and 0.05 % Tween 20 (AMRESCO) in dPBS (Gibco), normal goat serum (NGS) (Jackson Immuno Research) was diluted to 10% in the prepared blocking buffer, and blocking was performed for 90 min to minimize non-specific binding. The primary antibodies mentioned above were then immediately treated at the indicated optimal concentrations at 4 °C for 48 h for the primary reaction. Secondary antibodies, Alexa Fluor 594 anti-mouse (1:500) (Invitrogen) and Alexa Fluor 488 anti-rabbit (1:500) (Invitrogen) were also stained at 4 °C for 48 h for the secondary reaction. Finally, 4’, 6’-diamidine-2’-phenylindole dihydrochloride (DAPI) (1:1000) (Sigma) was treated for 90 min for nuclear labeling. All steps except the primary and secondary reaction were carried out at room temperature, and the washing process using dPBS was repeated 3 times for 10-20 min between all steps except the transition from blocking to the primary reaction. The remaining processes after the completion of the primary reaction was conducted in a light-blocked environment to prevent photobleaching. All fluorescence images were obtained using a laser scanning confocal microscope (ZEISS, LSM880), and the intensity of each fluorescence signal was quantitatively analyzed using the Image J program.

### Heart rate analysis

Cardiotoxicity evaluation was performed with hiPSC-CMs cultured for 7 days on the heart-on-a-chip and the 12 well plate by treating isoproterenol and nifedipine, which are representative drugs that cause changes in heart rate. Since the heart rate of hiPSC-CMs cultured in 12 well plates has a very large error range depending on each well, we selected some wells with heart rates close to the normal range of 60-90 bpm just before the cardiotoxicity test. For the cardiotoxicity evaluation, 1 ml of each of the media containing 0.1, 1, 5, 10, and 20 μM of isoproterenol and 0.1, 1, 5, 20, and 100 nM of nifedipine was prepared. For the chip, the drugs were treated by loading 200 μl of the prepared media into the two inlet wells and 100 μl of the prepared media into the two outlet wells, and for the plate, the drugs were treated by loading 600 μl of the same amount of media into the wells as was loaded into the chip. As soon as the drugs were administered to the chips and plates, the changes in heart rate were observed by performing video recordings at 1 h intervals for 24 h at 37°C using the inverted microscope equipped with the digital camera. Based on the acquired videos, the drug-induced changes in heart rate were manually counted, graphed, and analyzed.

### Video-based heart beating analysis (Motion tracking)

An open-source program developed for monitoring human movement, Kinovea, is originally used for sports analysis. Yet, we also applied to motion tracking of heartbeats to identify changes in speed (or intensity) and beating patterns depending on the drug, which could not be observed intuitively, except the heart rate. We observed the efficacy and toxicity of different types and concentrations of drugs through motion tracking based on the video results obtained after treatment with isoproterenol at concentrations of 0.1, 1, 5, 10, and 20 μM and nifedipine at concentrations of 0.1, 1, 5, 20, and 100 nM for 1 h in the previous cardiotoxicity evaluation. Each video was divided into 64 regions, and motion tracking was performed by selecting one marker in each region, and the results were graphed and analyzed by setting linear kinematics.

### Field potential measurement

The field potential was measured to evaluate whether electrophysiological changes of cardiomyocytes in 2D and 3D culture methods and in response to different drugs could be detected through the heart-on-a-chip. First, to check the difference in the culture method of cardiomyocytes, 3.2 x 10^5^ of hiPSC-CMs were seeded on the heart-on-a-chip integrated with cMEA and cultured with Cardiosight-S^®^ media that was added Cardiosight-S^®^ supplement diluted 100 times for 7 days. The conventional 60-electrode MEA (Multichannel systems) was coated with Matrigel^®^ hESC-Qualified, loaded 3.2 x 10^5^ of hiPSC-CMs using the pipette, and then incubated for 7 days under the same conditions. Then, 1 h prior to measurement, the same media minus the Cardiosight-S^®^ supplement was replaced for adaptive culture, and the field potential was measured after 1 h. On the other hand, to evaluate drug-dependent changes, hiPSC-CMs were cultured in the same manner on the heart-on-a-chip integrated with cMEA, the same media without the Cardiosight-S^®^ supplement was replaced, and the drug pre-treatment condition was measured after 1 h. Then, it was replaced with the same media to which 5 μM isoproterenol and 20 nM nifedipine were added, respectively, and the condition was measured 1 h after drug treatment. All measurements were taken using the Multichannel systems MEA2100-system and Cardio2D+ program (Multichannel systems). The analysis was based on the following specific outcomes: electrocardiogram waveform, heart rate (HR), T-wave amplitude, R-R wave interval mean (RR), and QT interval. In particular, for the QT interval, the results were corrected using QT interval correction formulas such as Van der Linde’s (QTcV=QT-0.087((60/HR)-1)) and Fridericia’s (QTcF=QT/(RR^1/3)).

### Total RNA-seq

Total RNA of hiPSC-CMS was isolated using Trizol reagent (Invitrogen). RNA quality was evaluated using an Agilent 2100 bioanalyzer with RNA 6000 Nano Chip (Agilent), and RNA quantification was carried out using an ND-2000 Spectrophotometer (Thermo). Libraries were prepared using an NEBNext Ultra II Directional RNA-Seq Kit (NEW ENGLAND BioLabs). Ribosomal RNA (rRNA) was removed using a RIBO COP rRNA depletion kit (LEXOGEN). The rRNA-depleted RNAs were used for the complementary DNA (cDNA) synthesis and shearing, following the manufacturer’s instructions. Indexing was conducted using an Illumina indexes1-12 (Illumina). The enrichment step was performed via PCR. Subsequently, libraries were checked using the Agilent 2100 bioanalyzer with DNA High Sensitivity Kit to assess the mean fragment size. Quantification was performed using a StepOne Real-Time PCR System (Life Technologies) with the library quantification kit. High-throughput sequencing was conducted as paired-end 100 sequencing using a NovaSeq 6000 (Illumina). A quality control of raw sequencing data was carried out through FastQC [111]. Adapter and low-quality reads (<Q20) were removed via FASTX_Trimmer [112] and BBMap [113]. Then, the trimmed reads were mapped to the reference genome through TopHat [114]. Gene expression levels of genes, isoforms and long non-coding RNAs (lncRNAs) were estimated using fragments per kb per million reads (FPKM) values via Cufflinks [115]. The FPKM values were normalized based on the Quantile normalization method Through EdgeR within R [116]. Data mining and graph visualization were performed using ExDEGA (Ebiogen).

### Statistical analysis

All data show mean ± SD. Statistical analysis were performed using unpaired t-test, one-way, and two-way ANOVA with Tukey’s multiple comparisons test through GraphPad Prism 9 for Windows. Differences between groups were considered statistically significant at *P*<0.05.

## Supporting Information

**Figure S1.**
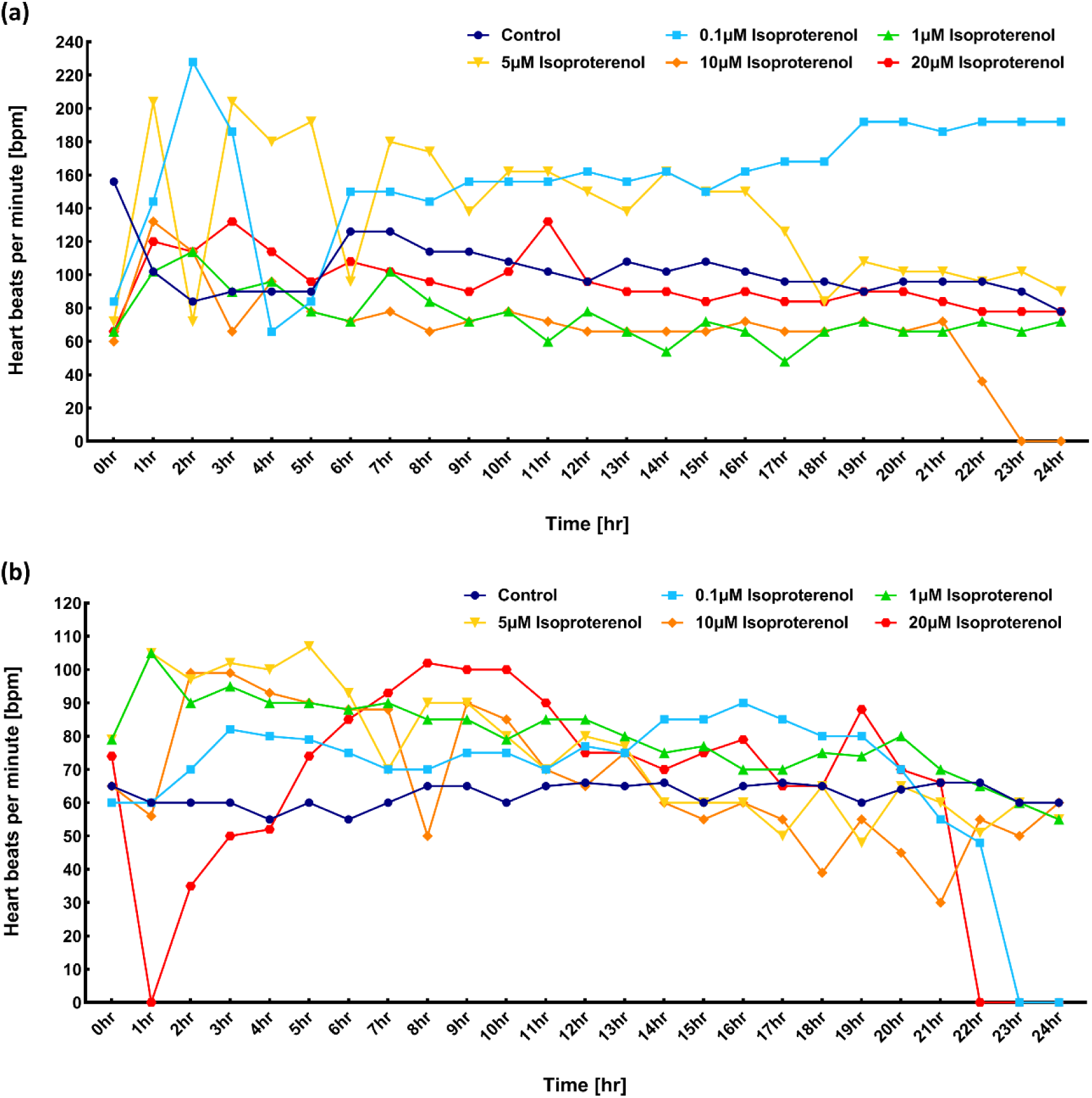
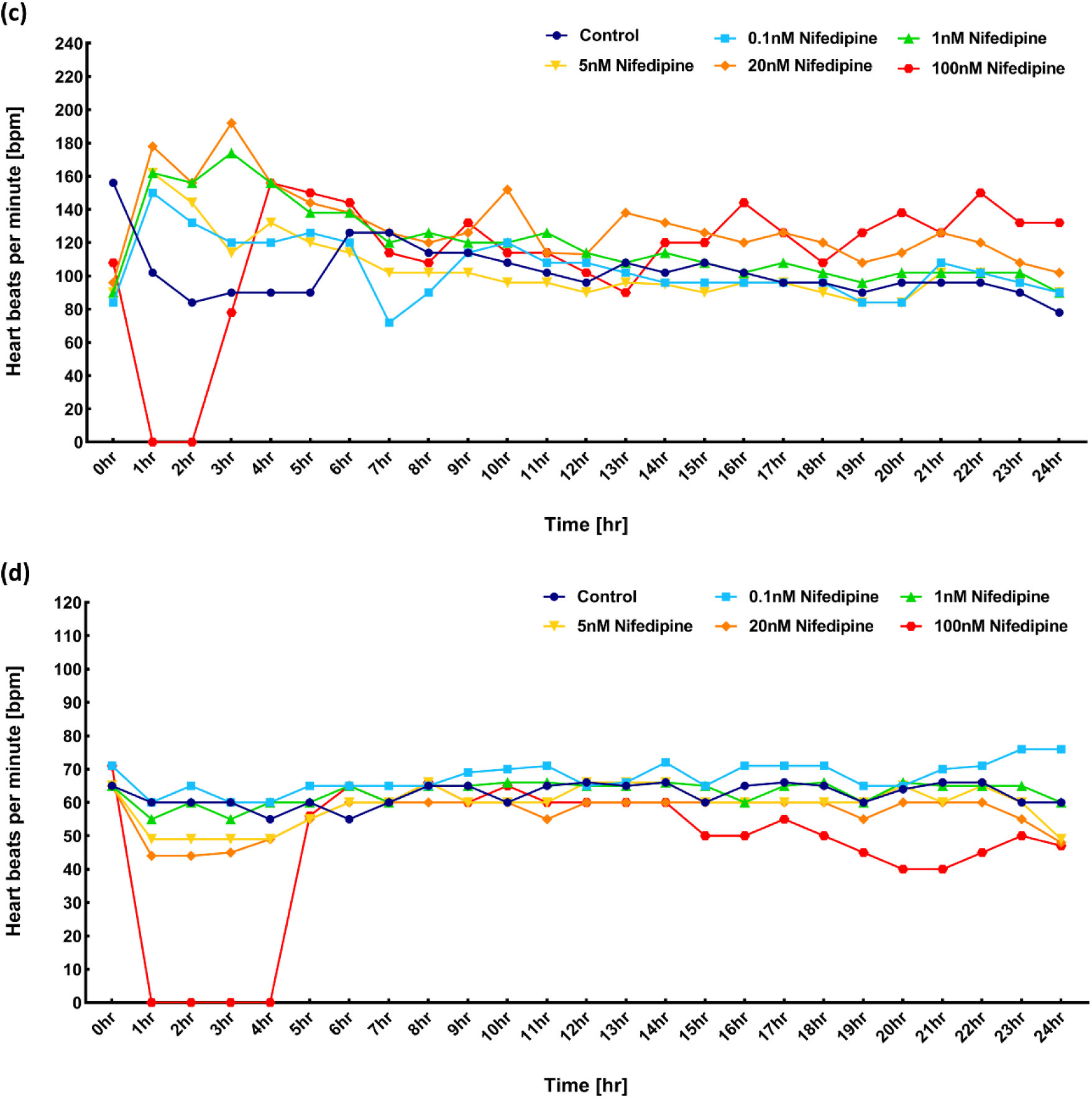
(a) Beating rate analysis on the 12 well plate with isoproterenol for 24 h. (b) Beating rate analysis on the heart-on-a-chip with isoproterenol for 24 h.(c) Beating rate analysis on the 12 well plate with nifedipine for 24 h. (d) Beating rate analysis on the heart-on-a-chip with nifedipine for 24 h.

**Figure S2.**
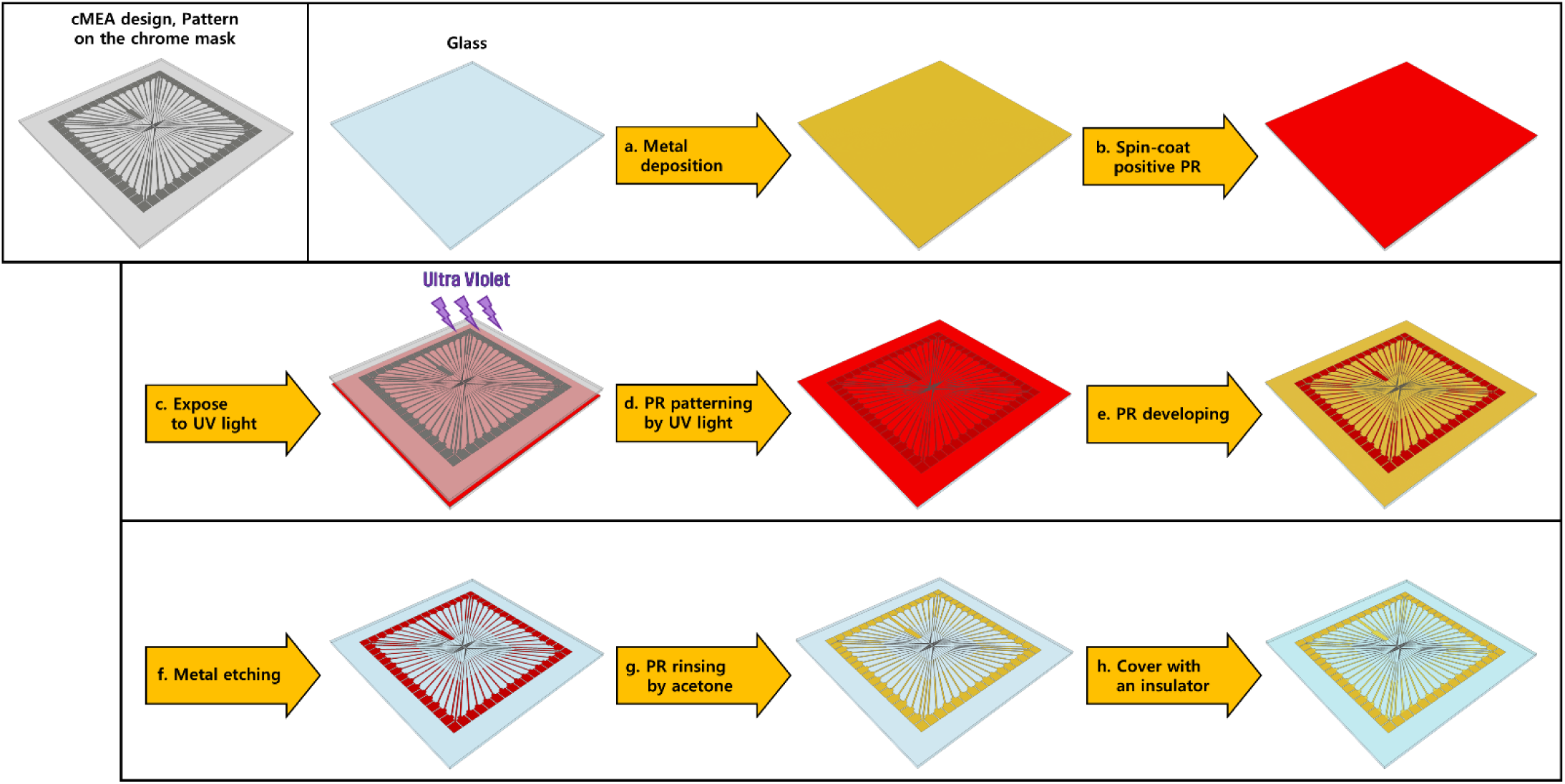
Fabrication protocol of the customized multi-electrode array (cMEA)

**Figure S3.**
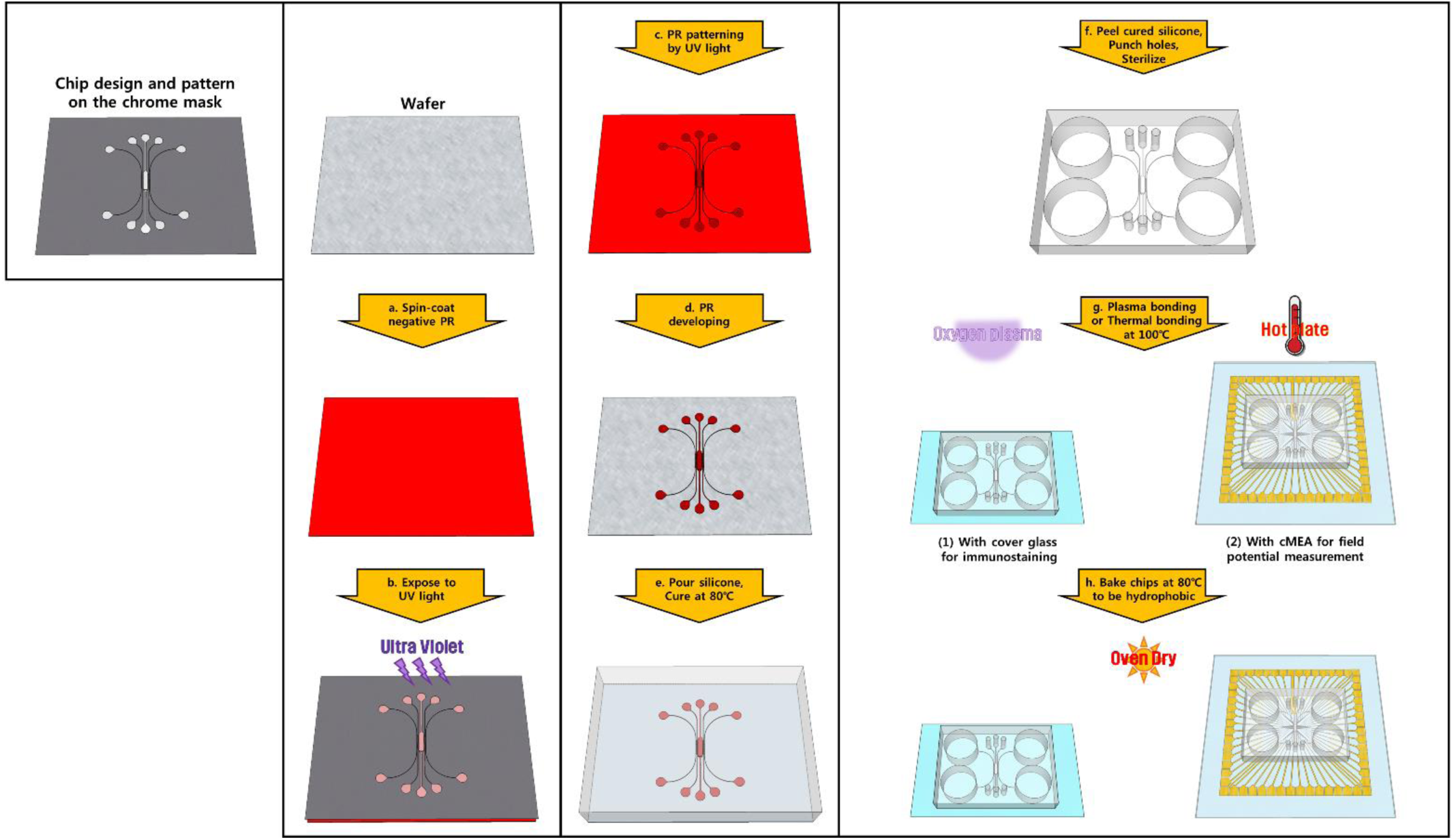
Fabrication protocol of the microfluidic platform

**Movie S1.** 3D modeling of cMEA-based heart-on-a-chip

**Movie S2.** Separation loading without micro-valve

**Movie S3.** Reversible bonding of microfluidics fabricated by PDMS and silicone adhesive mixture

**Movie S4.** Injection of hiPSC-CMs into the cMEA-based heart-on-a-chip

**Movie S5.** Beating of hiPSC-CMs on the cMEA-based heart-on-a-chip

**Movie S6.** Inconsistent heartbeats among each well of the 12 well plate

**Movie S7.** Consistent heartbeats among each heart-on-a-chip

**Movie S8.** Changes of heartbeats per minute on the 12 well plate for 7 DIV

**Movie S9.** Changes of heartbeats per minute on the heart-on-a-chip for 7 DIV

**Movie S10.** Dose-dependent beating rate analysis on the 12 well plate after isoproterenol treatment for 1 h

**Movie S11.** Dose-dependent beating rate analysis on the heart-on-a-chip after isoproterenol treatment for 1 h

**Movie S12.** Dose-dependent beating rate analysis on the 12 well plate after nifedipine treatment for 1 h

**Movie S13.** Dose-dependent beating rate analysis on the heart-on-a-chip after nifedipine treatment for 1 h

